# GPCR-Gαq Signaling Drives Adaptive Resistance to MEK Inhibition in BRAF-Mutant Melanoma

**DOI:** 10.1101/2025.11.18.688995

**Authors:** Jingyi Ke, Zi Wang, Zhengyang Guo, Guangshuo Ou, Wei Li

## Abstract

Oncogenic BRAFV600E mutations drive constitutive MAPK signaling across various cancers; however, resistance to MEK inhibitors such as trametinib undermines clinical benefits, and the mechanisms sustaining MAPK activity under treatment remain unclear. We developed a *lin-45(V627E) Caenorhabditis elegans* model to mimic oncogenic BRAF signaling and found that these animals tolerate ultra-high doses of trametinib. This survival is driven by an adaptive resistance bypass, mediated by Gαq-PLCβ-PKC signaling, which sustains MAPK activity despite MEK inhibition. Transcriptomic analyses of patient tumors and resistant melanoma cells reveal upregulation of the Gαq subunits and GPCR PAR2 upon acquisition of resistance. Functional inhibition of PAR2 or downstream Gαq signaling suppresses ERK reactivation, selectively impairs the viability of resistant cells. Furthermore, combined inhibition of MEK and Gαq-PLCβ-PKC signaling synergistically suppresses tumor growth and restores trametinib sensitivity in xenograft models. These findings uncover adaptive GPCR-Gαq-driven signaling rewiring as an evolutionarily conserved and therapeutically actionable vulnerability in MAPK-driven cancers.

## Introduction

Melanoma is a highly aggressive cancer with an increasing global incidence, and remains a leading cause of skin cancer-related mortality (Schadendorf et al. 2018; Centeno et al. 2023). A defining feature of melanoma is the constitutive activation of the MAPK pathway, most commonly driven by BRAFV600 mutations (Davies et al. 2002). Oncogenic BRAF triggers the RAF-MEK-ERK signaling cascade, promoting tumor growth and survival, providing a mechanistic basis for targeted therapies (Wan et al. 2004; Flaherty et al. 2012a; Flaherty et al. 2012b; Halle and Johnson 2021). As a result, BRAF and MEK inhibitors (BRAFi/MEKi) have significantly improved clinical outcomes, and combination BRAFi/MEKi therapy is now the standard of care for patients with BRAF-mutant melanoma (Dummer et al. 2018; Cohen and Sullivan 2019; Adashek et al. 2022). However, intrinsic and acquired resistance to these MAPK inhibitors remains a major challenge, limiting long-lasting therapeutic responses (Kakadia et al. 2018; Hartman et al. 2020). While genomic analyses of resistant tumors have identified diverse mechanisms of MAPK reactivation (Turajlic et al. 2014), the complexity and heterogeneity of these resistance pathways are not fully understood, in part due to the lack of genetically tractable models that can accurately replicate acquired BRAFV600E resistance.

Evolutionarily conserved model systems, such as *Caenorhabditis elegans*, provide powerful platforms to uncover the fundamental mechanisms of therapy resistance. *C. elegans*has proven to be a robust model for studying Ras-MAPK signaling (Sternberg and Han 1998), with highly conserved components including Ras/LET-60, RAF/LIN-45, MEK-2, and ERK/MPK-1 (Han and Sternberg 1990; Han et al. 1993; Lackner et al. 1994; Wu et al. 1995). MAPK signaling in worms regulates key developmental processes (Lackner and Kim 1998; Hsu et al. 2002; Sundaram 2013). Hyperactivation or inhibition of MAPK signaling produce quantifiable phenotypes, such as multivulva and vulvaless patterns, which can be used to evaluate drug responses (Hara and Han 1995; Bae et al. 2012; Ji et al. 2019; Gorgon et al. 2024). Complete loss of MAPK pathway activity results in sterility, highlighting its essential biological role (Church et al. 1995; Hajnal and Berset 2002). These features make *C. elegans*an ideal model for dissecting MAPK regulation and drug resistance.

Resistance to MAPK inhibitors in melanoma frequently arises through complex non-genetic mechanisms that restore MAPK pathway activity under therapeutic pressure (Rebecca et al. 2020). MAPK reactivation can be driven by compensatory signaling circuits, including ABL1/2-mediated MEK/ERK/MYC reactivation, PI3K-AKT-mTOR activation, and SRC family kinase signaling (Paraiso et al. 2011; Boshuizen et al. 2018; Tripathi et al. 2020). Notably, translational control mediated by the eIF4F complex acts as a central node sustaining oncogenic and survival programs under BRAF and MEK inhibition (Boussemart et al. 2014). In addition, metabolic reprogramming and post-translational signaling plasticity further sustain tumor survival, while receptor tyrosine kinases such as PDGFRB and EGFR contribute to bypass signaling (Nazarian et al. 2010; Gopal et al. 2014; Sun et al. 2014; Han et al. 2018). Bypass signaling, however, is not restricted to receptor tyrosine kinases. G protein-coupled receptors (GPCRs) have emerged as key regulators of oncogenic signaling and cancer progression (Dorsam and Gutkind 2007; O’Hayre et al. 2014; Stagg and Gutkind 2025). Among them, the protease-activated receptor PAR2 (*F2RL1*) promotes tumorigenesis in multiple cancer types (Kawaguchi et al. 2020; Bhansali et al. 2025; Dai et al. 2025). Gαq signaling also sustains MAPK activity in *GNAQ/GNA11*-mutant uveal melanoma (Chen et al. 2014; Maziarz et al. 2018; Hitchman et al. 2021; Ma et al. 2021) However, it remains unclear whether GPCR-Gαq signaling contributes to adaptive escape from MAPK inhibition in cutaneous BRAFV600E melanoma.

Here, we developed a*lin-45(V627E)C.elegans* model of acquired MEK inhibitor resistance, enabling systematic dissection of adaptive signaling that sustains MAPK reactivation under therapeutic pressure. Using this system, we identified a previously unrecognized GPCR-Gαq bypass pathway that sustains MAPK output despite MEK inhibition. This mechanism is conserved in human BRAFV600E melanoma, suggesting a potential strategy for overcoming resistance to MEK inhibitors.

## Results

### A*lin-45(V627E)C.elegans*Model Reveals Acquired Trametinib Resistance

To investigate mechanisms of acquired resistance to MEK inhibition in a genetically tractable in vivo system, we developed a *Caenorhabditiselegans*model harboring the oncogenic*lin-45(V627E)*mutation, orthologous to human BRAFV600E (Fig. S1A). Remarkably,*lin-45(V627E)*animals survive and reproduce at 100 μM trametinib, a concentration that completely abolishes fertility in wild-type worms, revealing an unanticipated and robust resistance phenotype (Fig. 1A). To assess the resistance profile, we exposed both wild-type and *lin-45(V627E)* animals to increasing concentrations of trametinib (1-100 μM) and measured fertility in the F1 generation (Fig. 1A). In wild-type animals, trametinib caused a dose-dependent loss of fertility, with complete sterility observed at 10 μM and 100 μM (Fig. 1B), consistent with MAPK pathway ablation (Sundaram 2013). In contrast, *lin-45(V627E)* mutants exhibited a nonlinear dose response: while 10 μM trametinib nearly abolished fertility, increasing the dose to 100 μM restored brood size to approximately 63% of untreated levels (Fig. 1B, Fig. S2C), indicating a capacity for resistance. This survival at ultra-high drug concentrations mirrors adaptive resistance observed in BRAFV600E melanoma cells following prolonged MEK inhibitor exposure.

**Figure 1.**
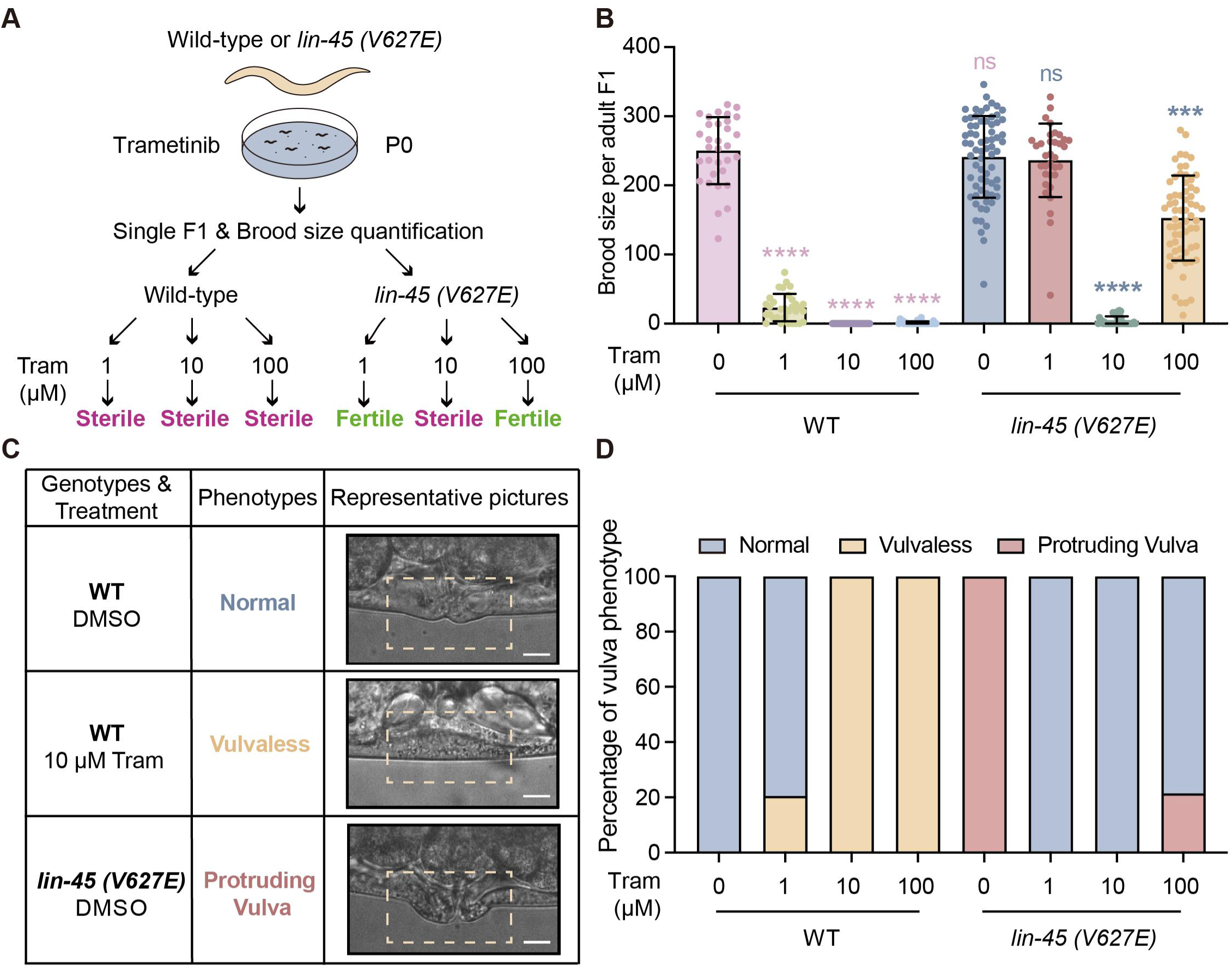
A *lin-45(V627E) C. elegans* model reveals acquired trametinib resistance under high concentration treatment. **(A)** Experimental workflow of brood size quantification in wild-type (WT) or *lin-45(V627E)* worms treated with DMSO or different concentrations of trametinib (Tram). **(B)** Dose-dependent effects of trametinib (Tram) on brood size of WT and *lin-45(V627E)* F1 animals. Statistical analysis was performed using Kruskal-Wallis test followed by Dunn’s multiple comparisons test. n > 30. *** P < 0.001, **** P < 0.0001; ns, not significant. **(C)** Representative bright-field images of vulva from WT and*lin-45(V627E)*mutants, treated with DMSO and 10 μM trametinib (Tram). n > 30. Scale bar, 100 μm. **(D)** Quantification of vulva phenotypes in WT and*lin-45(V627E)* mutants treated with DMSO or different concentrations of trametinib (Tram). n > 100.

Vulval phenotypes were tracked alongside fertility outcomes as secondary phenotypic readouts. Low and intermediate doses of trametinib suppressed the protruding vulva phenotype, whereas high-dose treatment (100 μM) selectively restored this phenotype in *lin-45(V627E)* mutants (Figs. 1C-D, Fig. S1B). These findings establish *lin-45(V627E)* worms as a genetically tractable in vivo model of acquired trametinib resistance, capable of tolerating MEK inhibitor doses that are otherwise lethal to wild-type animals.

### Transcriptomic and Genetic Analyses Implicate EGL-30/Gαq in the Resistant State

To identify pathways associated with acquired resistance, we performed RNA sequencing on trametinib-sensitive (10 μM) and trametinib-resistant (100 μM) *lin-45(V627E)* F1 animals. Principal component analysis revealed clear segregation between the two conditions, indicating a distinct transcriptional program associated with resistance (Fig. S2A). Differential expression analysis identified 443 genes significantly upregulated in resistant animals (fold change > 2, Padj < 0.05; Fig. 2A). Gene ontology analysis revealed strong enrichment for G-protein-coupled signaling, with *egl-30,*encoding the sole *C.elegans*Gαq ortholog, among the most significantly upregulated components (fold change = 2.3, Padj = 0.008; Figs. 2A-B, Fig. S2B). This transcriptional signature suggested that enhanced Gαq signaling correlates with the resistant state.

**Figure 2.**
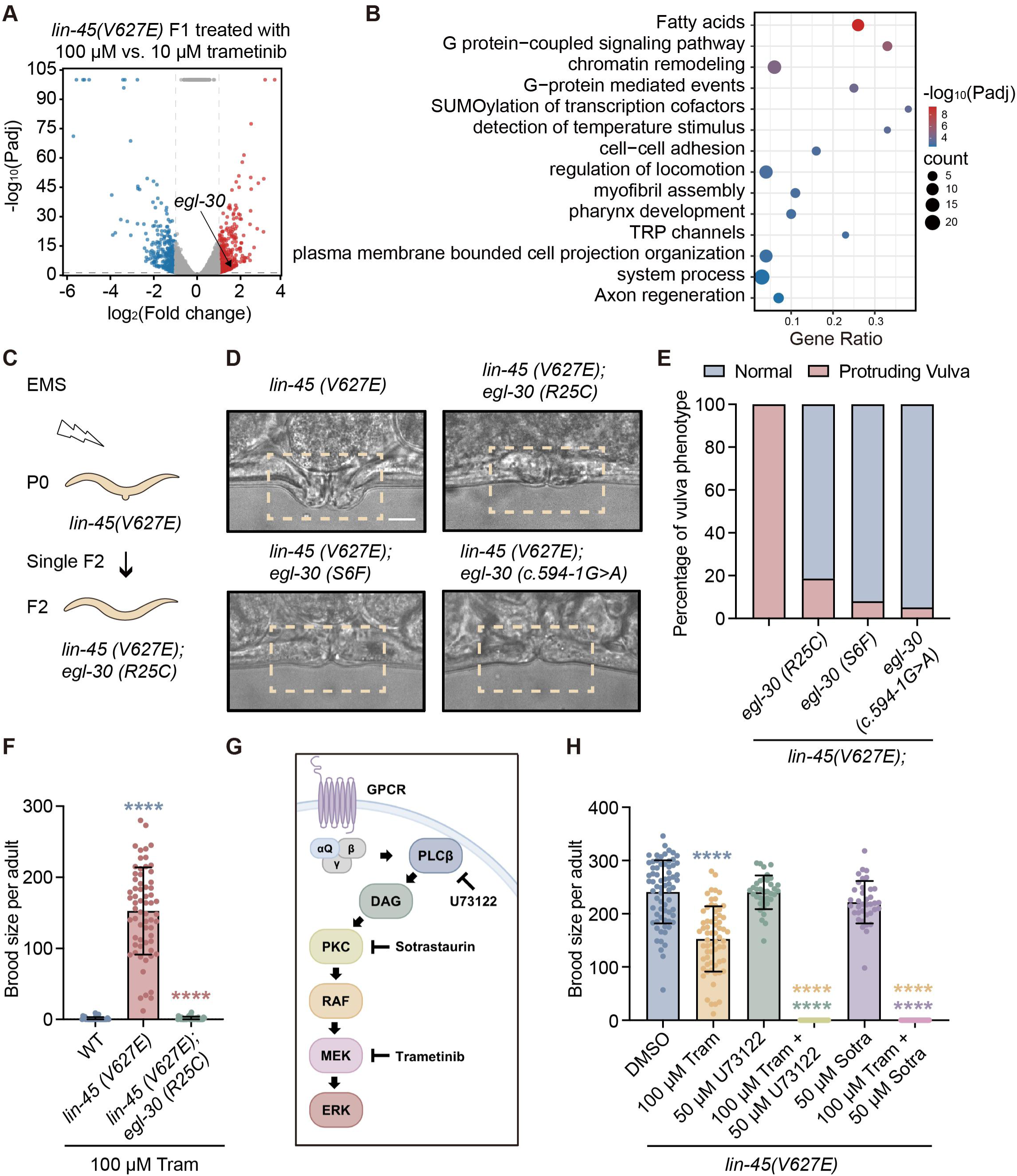
EGL-30/Gαq-PLCβ-PKC signaling is required for acquired trametinib resistance in*lin-45(V627E)*mutants. **(A)** Volcano plot of differentially expressed genes in*lin-45(V627E)* F1 animals treated with 100 μM trametinib relative to 10 μM trametinib. Upregulated genes are shown in red and downregulated genes in blue. Cutoffs: fold change > 2 and adjusted P value (Padj) < 0.05. n = 3 independent biological replicates per condition. **(B)** Gene Ontology (GO) enrichment analysis of upregulated genes in*lin-45(V627E)* F1 animals treated with 100 μM trametinib. Dot size represents gene counts, and color indicates enrichment significance. **(C)** Schematic overview of the EMS-based genetic suppressor screen in*lin-45(V627E)* animals to identify mutations restoring vulval development. **(D)** Representative images of vulval phenotypes of*lin-45(V627E)*,*lin-45(V627E); egl-30(R25C)*,*lin-45(V627E);egl-30(S6F)*, and*lin-45(V627E);egl-30(c.594-1G>A)* mutants. **(E)** Quantification of vulval phenotypes corresponding to quantifications in (D). n > 50 animals per genotype. **(F)** Brood size of WT,*lin-45(V627E)* and*lin-45(V627E); egl-30(R25C)* animals treated with 100 μM trametinib (Tram). Statistical analysis was performed using Kruskal-Wallis test followed by Dunn’s multiple comparisons test. n > 35 animals per genotype. ****P < 0.0001. **(G)** Schematic representation of the Gαq-PLCβ-PKC signaling pathway and the molecular targets of U73122, sotrastaurin, and trametinib. **(H)** Brood sizes of*lin-45(V627E)*F1 animals treated with DMSO, 100 μM trametinib (Tram), 50 μM U73122, 50 μM sotrastaurin (Sotra), or indicated drug combinations. Statistical analysis was performed using the Kruskal-Wallis test followed by Dunn’s multiple comparisons test. n > 35 animals per condition. ****P < 0.0001.

Independently, an unbiased forward genetic suppressor screen identified a recessive *egl-30(R25C)*allele that suppressed the protruding vulva phenotype of*lin-45(V627E)* mutants (Fig. 2C). Introduction of two additional *egl-30*loss-of-function alleles into the *lin-45(V627E)* background produced consistent suppression (Figs. 2D-E), reinforcing a correlative link between reduced Gαq activity and attenuation of *lin-45(V627E)*-driven phenotypes. Together, the transcriptomic and genetic data converge on EGL-30/Gαq as a key determinant of the trametinib-resistant state.

### Reduced EGL-30/Gαq Activity Compromises High-Dose Trametinib Resistance

To directly assess the role of EGL-30/Gαq in resistance, we examined fertility in *lin-45(V627E); egl-30(R25C)* double mutants. Although these animals exhibited reduced fertility compared to wild-type, they remained viable and produced measurable progeny (13 ± 7 per animal), enabling quantitative evaluation of drug response (Fig. S2E). Notably, reduction of EGL-30/Gαq activity sensitized animals to trametinib: while *lin-45(V627E)* single mutants regained fertility at 100 μM trametinib,*lin-45(V627E); egl-30(R25C)* mutants exhibited near-complete sterility under the same conditions (Fig. 2F). These results suggest that reduction of EGL-30/Gαq activity specifically disrupts high-dose trametinib resistance, indicating that Gαq signaling is functionally required for resistance state rather than merely correlated with it.

### Gαq Promotes Resistance via Its Canonical PLCβ-PKC Signaling Axis

Gαq classically signals through phospholipase Cβ (PLCβ), generating diacylglycerol and activating protein kinase C (PKC), which can directly stimulate MAPK pathway components (Hubbard and Hepler 2006). We therefore hypothesized that an EGL-30/Gαq-PLCβ-PKC axis enables MAPK reactivation under MEK blockade (Fig. 2G). To test this,*lin-45(V627E)*animals were treated with the PLCβ inhibitor U73122 or the PKC inhibitor sotrastaurin, alone or in combination with trametinib. Neither inhibitor alone substantially affected fertility, indicating minimal baseline toxicity (Fig. 2H, Fig. S2D). In contrast, co-treatment with 100 μM trametinib and either inhibitor resulted in complete sterility, phenocopying genetic suppression of *egl-30* (Fig. 2H, Fig. S2D). These results show that activities of Gαq’s downstream effectors are required for high-dose trametinib resistance, implicating PLCβ and PKC as key mediators of MAPK pathway rebound in vivo.

### Gαq-PLCβ-PKC Inhibition Synergizes with MEK Blockade in BRAFV600E Melanoma Cells and Xenografts

To determine whether this resistance mechanism is conserved in human cancer, we examined BRAFV600E A375 melanoma cells. shRNA-mediated knockdown of the Gαq subunits *GNAQ* and *GNA11* (Fig. 3A) significantly lowered the IC_50_ of trametinib (Figs. 3B-C) and reduced ERK phosphorylation (Fig. 3D), indicating that Gαq activity contributes to MAPK signaling under MEK inhibition. Pharmacological inhibition of PLCβ or PKC in combination with trametinib also produced robust synergistic suppression of cell proliferation, with combination index values <0.3 across most dose combinations (Figs. 3E-F). Correspondingly, ERK phosphorylation was more effectively suppressed by combination therapy than by either agent alone (Fig. 3G). In parallel, both genetic knockdown and pharmacological inhibition of Gαq signaling markedly impaired clonogenic growth in A375 cells (Figs. S3A-C), reinforcing the contribution of Gαq to proliferative capacity under targeted therapy pressure. These findings recapitulate the genetic interactions observed in *C.elegans*, demonstrating evolutionary conservation of adaptive resistance mechanism.

**Figure 3.**
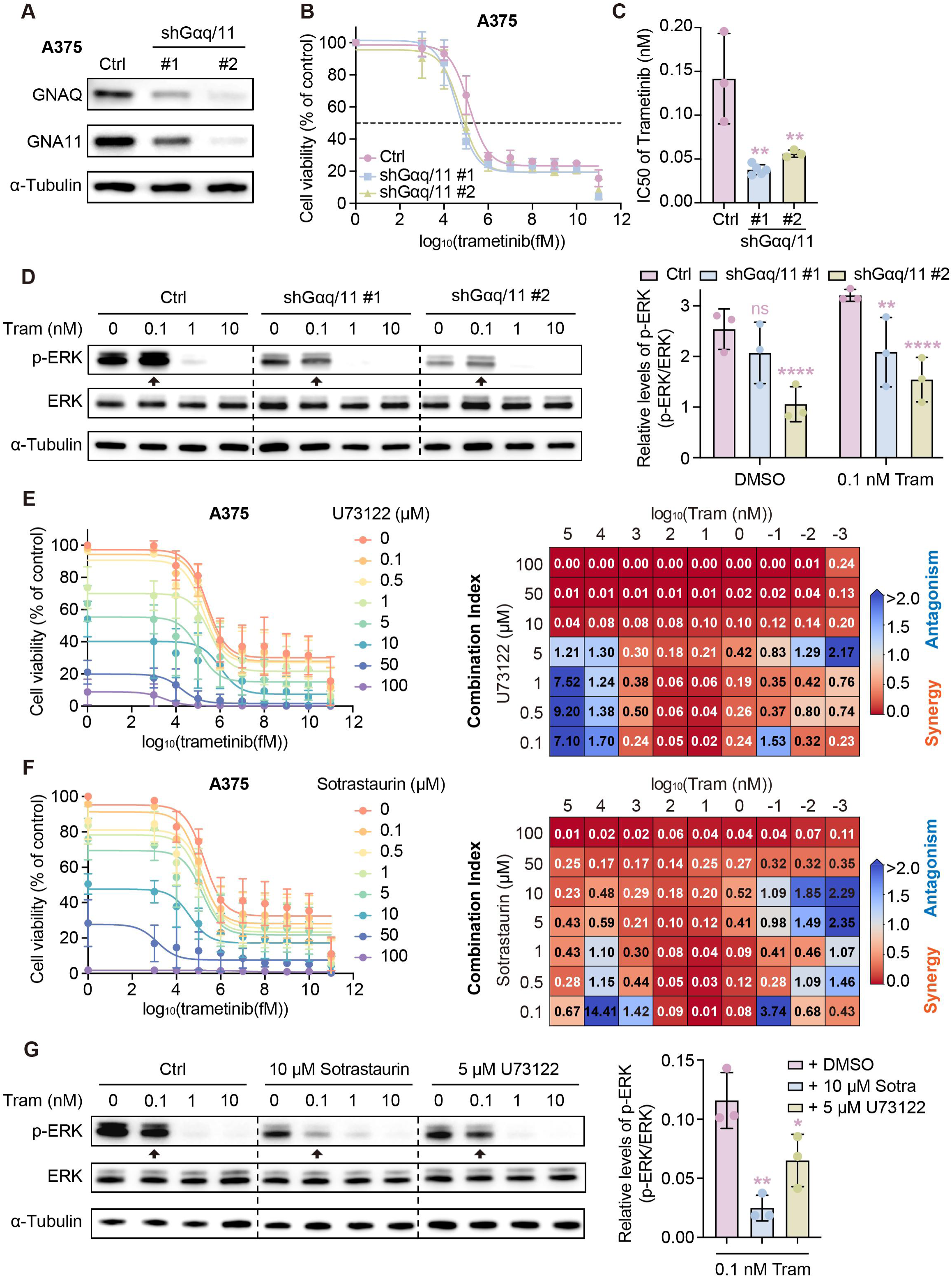
Synergistic suppression of A375 cell proliferation by combined Gαq/11-PLCβ-PKC blockade and MEK inhibition. **(A)** Knockdown efficiency of *GNAQ* and *GNA11* in A375 cells transduced with shRNA targeting both genes (shGαq/11), assessed by immunoblotting. **(B)** A375 cells transduced with control vector (Ctrl) or shGαq/11 were treated with vehicle or trametinib for 72 h. Cell viability was measured using CellTiter-Glo, and dose-response curves are shown (mean ± SD, n = 3 independent experiments). **(C)** IC_50_ values of trametinib in Ctrl- and shGαq/11-transduced A375 cells. n = 3 independent experiments. **P < 0.01; one-way ANOVA with Tukey’s multiple comparisons test. **(D)** Western blot analysis of indicated proteins in Ctrl- or shGαq/11-transduced A375 cells treated with DMSO or trametinib at the indicated concentrations for 24 h. Representative blots (left; n = 3 independent experiments) and quantification of p-ERK levels (right) are shown. **P < 0.01, ****P < 0.0001; ns, not significant; two-way ANOVA with Tukey’s multiple comparisons test. **(E)** Dose-response curves of cell viability (left) and Chou-Talalay combination index (CI) plots (right) for A375 cells treated with U73122 in combination with trametinib at the indicated concentrations for 72 h (mean ± SD, n = 3). CI < 0.3 indicates strong synergy (red), 0.3-0.9 synergy, 0.9-1.1 additive effects, and ≥ 1.1 antagonism (blue). **(F)** Dose-response curves of cell viability (left) and CI plots (right) for A375 cells treated with sotrastaurin in combination with trametinib at the indicated concentrations for 72 h (mean ± SD, n = 3). **(G)** Western blot analysis of indicated proteins in A375 cells treated with DMSO, 10 μM sotrastaurin, or 5 μM U73122, alone or in combination with trametinib at the indicated concentrations for 24 h. Representative blots (left; n = 3 independent experiments) and quantification of p-ERK levels (right) are shown. *P < 0.05, **P < 0.01; one-way ANOVA with Tukey’s multiple comparisons test.

We next evaluated the therapeutic relevance in vivo using A375 xenografts in the mouse model. While trametinib, U73122, or sotrastaurin monotherapy delayed tumor growth, combination treatments produced substantially enhanced tumor suppression (Figs. 4A-F). Trametinib-sotrastaurin regimens caused no overt toxicity, while U73122-containing combinations induced modest, dose-dependent weight loss without additional adverse effects (Figs. S4A-B). The animals remained otherwise healthy, indicating dose-dependent systemic effects. These data establish that pharmacological disruption of Gαq-PLCβ-PKC signaling enhances MEK inhibitor efficacy in vivo.

**Figure 4.**
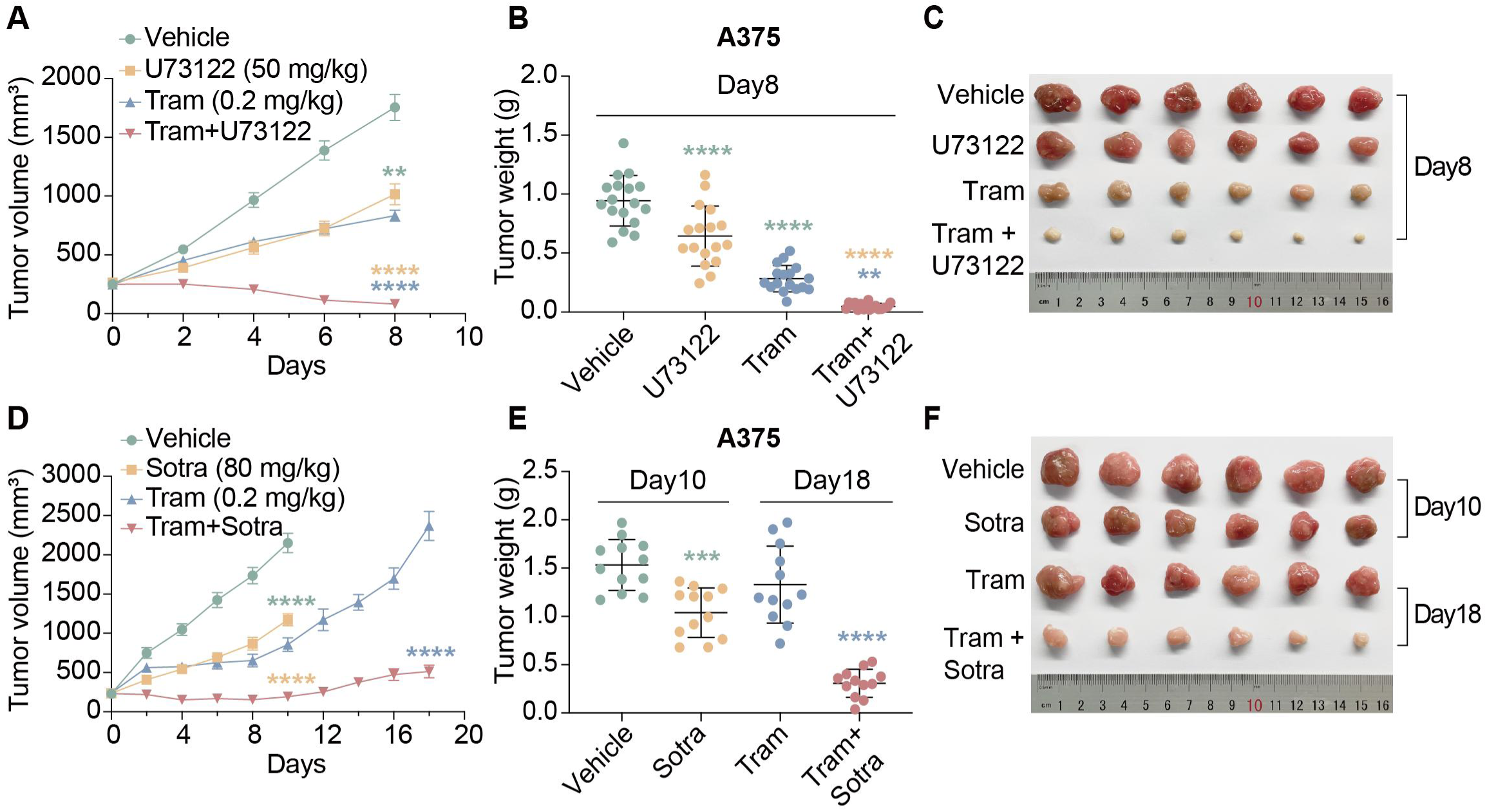
PLCβ-PKC blockade enhances MEK inhibitor efficacy in A375 xenograft models. **(A-C)** Parental A375 xenografts were randomized into treatment groups and administered vehicle, trametinib (0.2 mg/kg), U73122 (50 mg/kg) or their combination when mean tumor volume reached ∼250 mm³ (n = 17 per group). Tumor growth curves (A), endpoint tumor weights (B), and representative excised tumors (C) are shown. **(D-F)** Parental A375 xenografts treated with vehicle, trametinib (0.2 mg/kg), sotrastaurin (Sotra; 80 mg/kg), or their combination (n = 12 per group). Tumor growth curves (D), endpoint tumor weights (E), and representative excised tumors (F) are shown. Data represent mean ± SEM. Treatments were initiated when mean tumor volume reached ∼250 mm³. Statistical significance was determined by two-way ANOVA for tumor growth and one-way ANOVA for tumor weight, followed by Tukey’s multiple comparisons test, except for the comparison in (E) between the two indicated time-matched groups, which was analyzed using an unpaired two-tailed *t*-test. **P < 0.01, ***P < 0.001, ****P < 0.0001.

### Gαq-PLCβ-PKC Signaling Is Required to Maintain Acquired Trametinib Resistance

We next investigated whether Gαq signaling is required for maintaining an established resistant state in cells that have already developed resistance to MEK inhibition. To address this, we generated trametinib-resistant A375 cells (A375-TR) using an established protocol of chronic drug exposure (Villanueva et al. 2010). A375-TR cells exhibited a >500-fold increase in trametinib IC_50_, sustained ERK activation, and enhanced clonogenic capacity under high-dose treatment (Figs. 5A-C, Fig. S5A). Genetic suppression of *GNAQ* and *GNA11* markedly reduced ERK phosphorylation and restored trametinib sensitivity in A375-TR cells (Figs. 5D-G). Likewise, pharmacological inhibition of PLCβ or PKC selectively impaired viability of resistant cells and strongly synergized with trametinib (Figs. 5H-J). Consistently, both genetic knockdown and pharmacological inhibition of Gαq signaling substantially suppressed clonogenic outgrowth of A375-TR cells, indicating loss of long-term proliferative capacity under targeted therapy pressure (Figs. S5B-D). These data demonstrate that persistent Gαq-PLCβ-PKC signaling is essential to sustain MAPK activity and maintain the resistant state.

**Figure 5.**
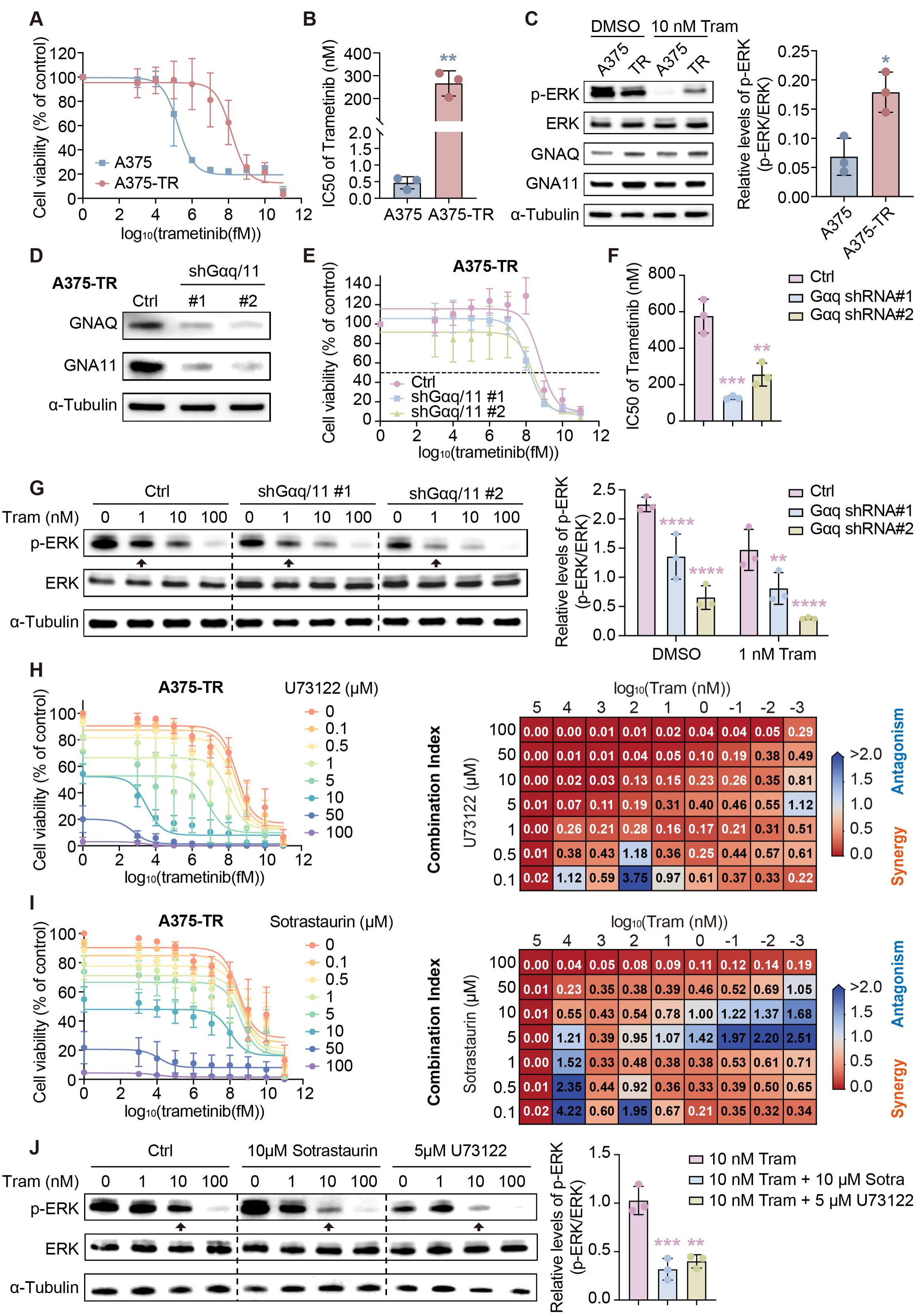
Synergistic suppression of trametinib-resistant A375 (A375-TR) cell proliferation by dual Gαq/11-PLCβ-PKC and MEK blockade. **(A)** Parental A375 and trametinib-resistant A375 (A375-TR) cells were treated with vehicle or trametinib for 72 h. Cell viability was measured using CellTiter-Glo, and dose-response curves are shown (mean ± SD, n = 3 independent experiments). **(B)** IC_50_ values of trametinib in A375 and A375-TR cells. n = 3, ** P < 0.01; unpaired two-tailed *t*-test. **(C)** Western blot analysis of indicated proteins in A375 and A375-TR cells treated with DMSO or 10nM trametinib for 24 h. Representative blots (left) (n = 3 independent experiments) and quantification of p-ERK (right) are shown, * P < 0.1; unpaired two-tailed *t*-test. **(D)** Knockdown efficiency of *GNAQ* and *GNA11* in A375 cells transduced with shGαq/11, assessed by immunoblotting. **(E)** Ctrl- and shGαq/11-transduced A375-TR cells were treated with vehicle or trametinib for 72 h. Cell viability was measured using CellTiter-Glo, and dose-response curves are shown (mean ± SD, n = 3 independent experiments). **(F)** IC_50_ values of trametinib in Ctrl- and shGαq/11-transduced A375-TR cells. n = 3 independent experiments. **P < 0.01, ***P < 0.001; one-way ANOVA with Tukey’s multiple comparisons test. **(G)** Western blot analysis of indicated proteins in Ctrl- or shGαq/11-transduced A375-TR cells treated with DMSO or trametinib at the indicated concentrations for 24 h. Representative blots (left; n = 3 independent experiments) and quantification of p-ERK levels (right) are shown. **P < 0.01, ****P < 0.0001; two-way ANOVA with Tukey’s multiple comparisons test. **(H)** Dose-response curves of cell viability (left) and CI plots (right) for A375-TR cells treated with U73122 in combination with trametinib at the indicated concentrations for 72 h (mean ± SD, n = 3). **(I)** Dose-response curves of cell viability (left) and CI plots (right) for A375-TR cells treated with sotrastaurin in combination with trametinib at the indicated concentrations for 72 h (mean ± SD, n = 3). **(J)** Western blot analysis of indicated proteins in A375-TR cells treated with DMSO, 10 μM sotrastaurin, or 5 μM U73122, alone or in combination with trametinib at the indicated concentrations for 24 h. Representative blots (left; n = 3 independent experiments) and quantification of p-ERK levels (right) are shown. **P < 0.01, ***P < 0.001; one-way ANOVA with Tukey’s multiple comparisons test.

We examined whether this dependency operates in vivo. Xenografts established from A375-TR cells were refractory to trametinib treatment, confirming their resistant state. In contrast, inhibition of PLCβ or PKC alone modestly suppressed tumor growth, while combination treatment with trametinib led to marked tumor regression (Figs. 6A-F). As expected, trametinib-sotrastaurin treatment caused no detectable adverse effects. We reduced the dosing frequency of the PLCβ inhibitor U73122 from three to two administrations per week, preserving robust antitumor efficacy while significantly alleviating treatment-associated body weight loss, indicating an improved therapeutic window (Figs. S6A-B).

**Figure 6.**
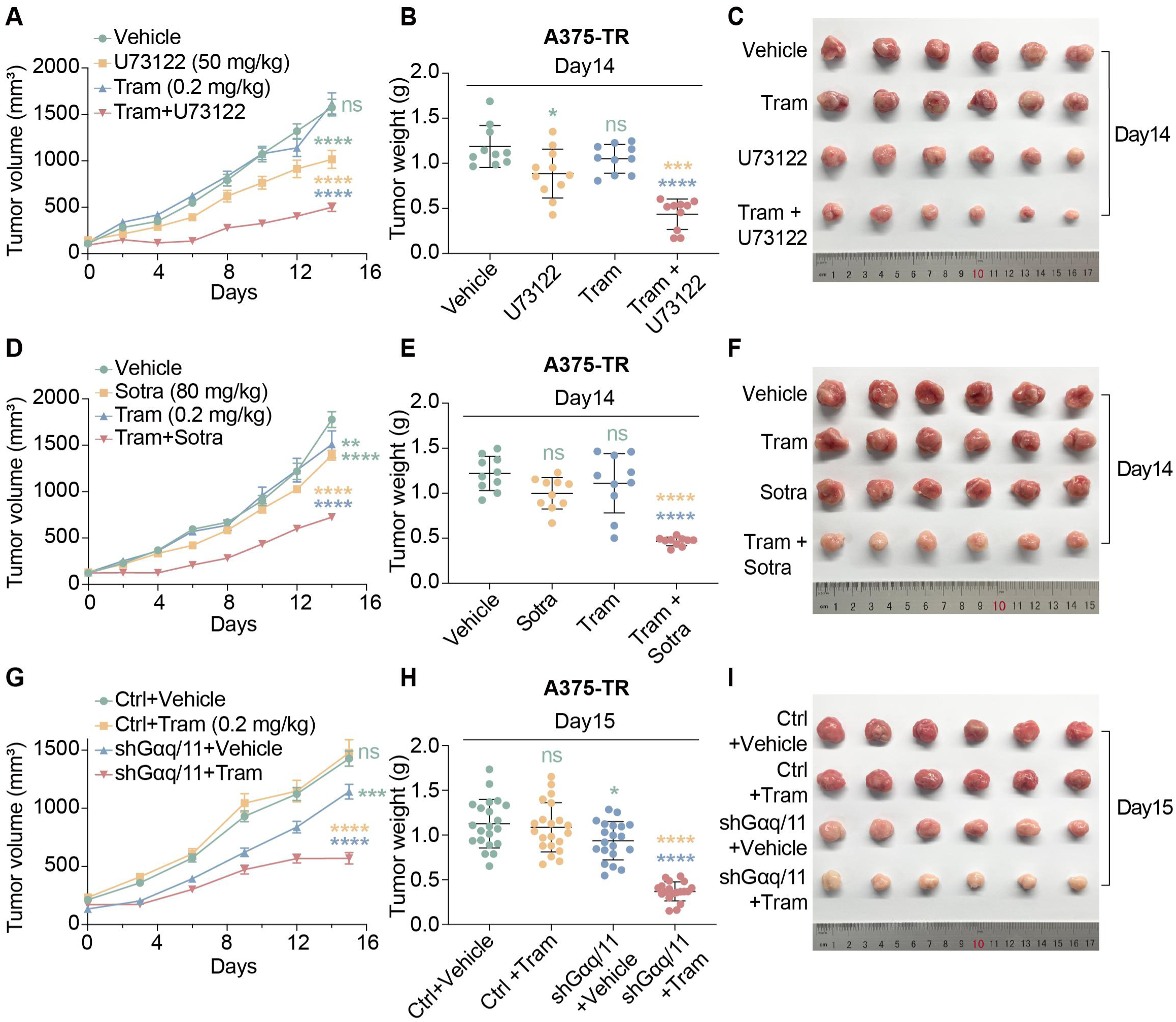
PLCβ-PKC blockade enhances MEK inhibitor efficacy in A375-TR xenograft models. (A-C) Trametinib-resistant A375-TR xenografts treated with vehicle, trametinib (0.2 mg/kg), U73122 (50 mg/kg), or their combination (n = 10 per group). Tumor growth curves (G), endpoint tumor weights (H), and representative excised tumors (I) are shown. **(D-F)** A375-TR xenografts treated with vehicle, trametinib (0.2 mg/kg), sotrastaurin (Sotra; 80 mg/kg), or their combination (n = 10 per group). Tumor growth curves (J), endpoint tumor weights (K), and representative excised tumors (L) are shown. **(G-I)** A375-TR xenografts transduced with control vector (Ctrl) or shGαq/11 and treated with vehicle or trametinib (0.2 mg/kg) (n = 18 mice per group). Tumor growth curves (M), endpoint tumor weights (N), and representative excised tumors (O) are shown. Data represent mean ± SEM. Treatments were initiated when mean tumor volume reached ∼250 mm³. Statistical significance was determined by two-way ANOVA for tumor growth and one-way ANOVA for tumor weight, followed by Tukey’s multiple comparisons test. *P < 0.1, ***P < 0.001, ****P < 0.0001; ns, not significant.

To genetically validate the requirement for Gαq signaling in an established resistant state in vivo, we generated xenografts from *GNAQ/11*-depleted A375-TR cells. These tumors exhibited significantly impaired growth compared with empty vector controls. Notably, trametinib treatment effectively suppressed growth of *GNAQ/11*-depleted xenografts, demonstrating that genetic disruption of Gαq signaling restores MEK inhibitor sensitivity in resistant tumors (Figs. 6G-I, Fig. S6C). Collectively, Gαq-PLCβ-PKC signaling is functionally required to maintain acquired trametinib resistance in vivo, and disruption of this pathway restores therapeutic vulnerability to MEK blockade.

### Adaptive Upregulation of PAR2-Gαq Signaling in Trametinib-Resistant Melanoma

To investigate the clinical relevance of Gαq signaling in MAPK inhibitor resistance, we analyzed RNA-seq datasets from melanoma patients both before treatment and after acquisition of resistance to BRAFi and/or MEKi therapy (Gopal et al. 2014; Long et al. 2014; Rizos et al. 2014; Sun et al. 2014; Wagle et al. 2014; Tirosh et al. 2016). Despite evident inter-patient heterogeneity, a substantial subset of resistant tumors exhibited increased expression of Gαq paralogs *GNAQ*, *GNA11*, and the GPCR *F2RL1* (PAR2) relative to matched pretreatment samples (Fig. 7A, Fig. S7A), suggesting convergent activation of PAR-Gαq signaling during therapeutic escape.

**Figure 7.**
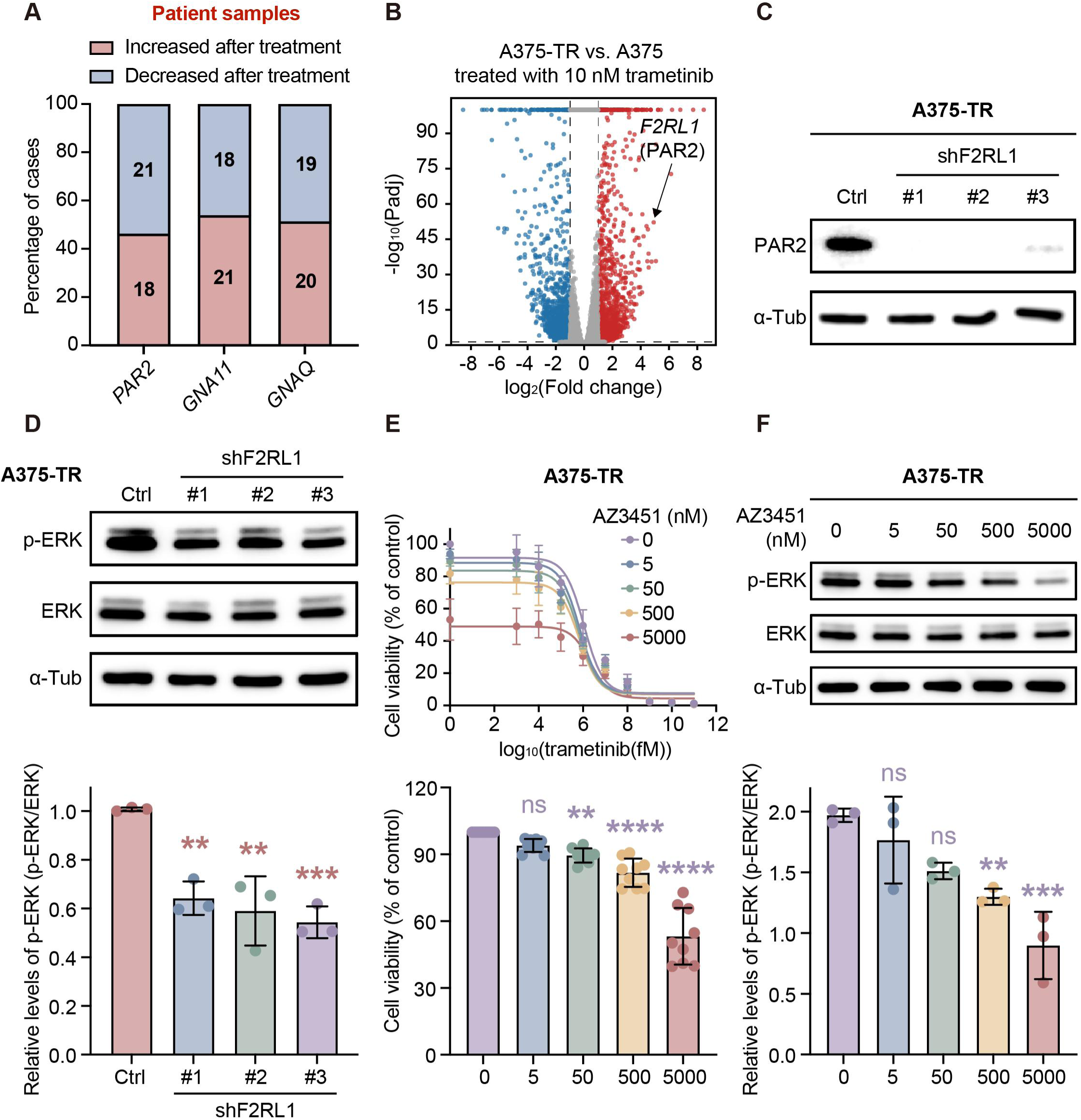
Upregulation of *F2RL1* (PAR2) sustains ERK signaling in trametinib-resistant melanoma cells. **(A)** Integrated analysis of primary human paired melanoma datasets (RNA-seq: 50535, 77940, 61992; microarray: 50509) were analyzed for *F2RL1*(PAR2), *GNAQ*, and *GNA11* mRNA levels in tumors pre-/post-treatment with BRAFi, MEKi, or combined BRAFi/MEKi. Numbers in each column indicate the number of cases. **(B)** Volcano plot of differentially expressed genes in trametinib-resistant A375 cells (A375-TR) treated with 10 nM trametinib relative to parental A375 cells treated under the same conditions. Upregulated genes are shown in red and downregulated genes in blue. Cutoffs: fold change > 2 and adjusted P value (Padj) < 0.05. n = 3 independent biological replicates per condition. **(C)** Knockdown efficiency of *F2RL1* in A375-TR cells transduced with shRNA targeting *F2RL1*(shF2RL1), assessed by immunoblotting. **(D)** Western blot analysis of indicated proteins in Ctrl- or shF2RL1-transduced A375-TR cells treated with 10 nM trametinib for 24 h. Representative blots (top; n = 3 independent experiments) and quantification of p-ERK levels (bottom) are shown. **P < 0.01, ***P < 0.001; one-way ANOVA with Tukey’s multiple comparisons test. **(E)** Dose-response curves of cell viability in A375-TR cells treated with AZ3451 in combination with trametinib at the indicated concentrations for 96 h (top, mean ± SD, n = 8). Cell viability of A375-TR cells treated with AZ3451 alone in the absence of trametinib for 96 h (bottom, n = 8). **P < 0.01, ****P < 0.000; ns, not significant; one-way ANOVA with Tukey’s multiple comparisons test. **(F)** Western blot analysis of indicated proteins in A375-TR cells treated with DMSO or AZ3451 at the indicated concentrations for 96 h. Representative blots (top; n = 3 independent experiments) and quantification of p-ERK levels (bottom) are shown. **P < 0.01, ***P < 0.001; ns, not significant; one-way ANOVA with Tukey’s multiple comparisons test.

To validate this transcriptional signature experimentally, we performed RNA-seq on trametinib-resistant A375 cells. Consistent with patient data, the GPCR *F2RL1*was robustly upregulated in resistant cells compared to parental controls (Fig. 7B). These findings highlight adaptive induction of a GPCR-Gαq signaling module as a recurrent transcriptional feature of trametinib resistance, linking clinical observations with the genetic and pharmacological mechanisms uncovered in this study.

### The GPCR PAR2 Drives ERK Reactivation and Sustains Trametinib Resistance

We next tested whether PAR2 directly contributes to MAPK reactivation in resistant melanoma cells. To this end, we generated three independent RNAi constructs targeting PAR2 (*F2RL1*), all of which effectively reduced PAR2 level (Fig. 7C). PAR2 depletion markedly diminished ERK phosphorylation in trametinib-resistant cells without inducing overt cytotoxicity (Fig. 7D), indicating that PAR2 is essential for sustaining MAPK signaling under MEK inhibition.

We next explored whether pharmacological inhibition of PAR2 could suppress the resistant phenotype. Treatment of A375-TR cells with the selective PAR2 antagonist AZ3451 significantly impaired cell viability and strongly reduced ERK phosphorylation (Figs. 7E-F). Notably, PAR2 inhibition phenocopied suppression of the downstream Gαq-PLCβ-PKC axis, positioning PAR2 as an upstream driver of adaptive MAPK reactivation. These findings collectively establish PAR2 as a critical upstream activator of Gαq-dependent MAPK rebound, revealing a non-genetic GPCR-mediated mechanism through which BRAFV600E melanoma cells adapt to MEK inhibition.

## Discussion

This study uncovers a pivotal role for GPCR-Gαq-PLCβ-PKC signaling in mediating adaptive resistance to trametinib in BRAFV600E melanoma, offering insights into how MAPK pathway output is sustained despite MEK inhibition. The ability of *lin-45(V627E)*worms to remain fertile under MEK inhibition—a condition that would otherwise induce sterility in wild-type worms—allowed the unbiased discovery of this resistance mechanism (Figs. 1A-B). Importantly, this finding was validated in human melanoma cell lines (Fig. 3 and Fig. 5) and xenografts (Figs. 4A-F), demonstrating that the cross-species approach used here—starting with mechanistic discovery in worms followed by translational validation in mammalian systems—can effectively uncover conserved resistance pathways that might be obscured in more complex tumor systems.

The therapeutic relevance of these findings is further supported by our in vivo data. Trametinib-resistant xenografts were resensitized to therapy when Gαq-PLCβ-PKC signaling was inhibited (Figs. 6A-F). Combined MEK and PKC inhibition induced marked tumor suppression with limited systemic toxicity, indicating a favorable therapeutic window (Fig. S6B). This underscores the potential of co-targeting MEK and Gαq-PKC signaling as an effective strategy to overcome acquired resistance in MAPK-driven cancers. One of the key insights from this study is the identification of PAR2, a Gαq-coupled GPCR (Adams et al. 2011), as a major contributor to resistance. PAR2 upregulation was consistently observed in both trametinib-resistant melanoma cell lines and clinical tumor samples (Figs. 7A-B), and its inhibition—either genetically or pharmacologically—resulted in significant suppression of ERK phosphorylation and cell viability (Figs. 7D-F). These findings position PAR2 as an upstream activator of Gαq-dependent MAPK reactivation, revealing a mechanism by which BRAFV600E melanoma cells bypass MEK inhibition. Importantly, this mechanism operates without the need for new genetic mutations, emphasizing the role of non-genetic signaling plasticity in therapeutic escape.

Importantly, this study highlights the power of model organisms — particularly *C. elegans*—as genetically tractable platforms for uncovering conserved mechanisms of therapeutic resistance. The capacity to perform unbiased, large-scale genetic and pharmacological screens in *C. elegans* enables rapid identification of resistance pathways and druggable vulnerabilities that may remain obscured in mammalian systems. Moreover, the ease of generating resistant mutants and mapping genetic interactions allows systematic dissection of complex adaptive signaling networks. Together, these features position *C. elegans* as a powerful discovery engine for revealing evolutionarily conserved resistance mechanisms and informing therapeutic strategies in human cancer.

Notably, analysis of the GSE245262 dataset, which profiles NRASQ61L melanoma cells rendered resistant to combined BRAF and MEK inhibition, revealed a striking upregulation of the *F2RL1* (PAR2) upon acquisition of resistance (approximately 16-fold) (Dinter et al. 2024). This response closely parallels our observations in BRAFV600E melanoma, suggesting that induction of *F2RL1* may represent a conserved adaptive response to MAPK pathway inhibition in MAPK-hyperactivated melanoma. Together, these findings support the idea that GPCR-Gαq signaling constitutes a shared mechanism engaged upon BRAFi/MEKi treatment across genetically distinct melanoma subtypes.

Beyond melanoma, our findings suggest broader implications for MAPK-driven malignancies. PAR2 has been implicated in tumor progression in multiple cancer types, including colorectal and breast cancer (Chaudhary and Kim 2021), while Gαq signaling is a well-established oncogenic driver in uveal melanoma through recurrent mutations in *GNAQ* and *GNA11* (Chen et al. 2014; Hitchman et al. 2021). Together, these observations raise the possibility that GPCR-Gαq-dependent signaling may represent a generalizable mechanism of adaptive MAPK reactivation. Thus, targeting this axis could hold therapeutic potential across diverse cancers that harbor MAPK pathway alterations or exploit similar adaptive resistance programs.

However, several key questions remain. Future studies will need to investigate the endogenous PAR2 ligands that operate in resistant tumors and explore the contributions of the tumor microenvironment in modulating GPCR-driven resistance pathways. Moreover, functional redundancy among other GPCRs may provide alternative routes to Gαq activation, warranting systematic exploration of alternative resistance networks (Adams et al. 2011).

In conclusion, our study identifies GPCR-Gαq-PLCβ-PKC signaling as a conserved and essential mechanism of adaptive resistance to MEK inhibition, revealing a therapeutically actionable vulnerability in BRAFV600E melanoma. By leveraging *C. elegans* as a genetically tractable in vivo discovery platform, we establish a cross-species strategy that enables systematic identification of conserved resistance circuits that may remain obscured in mammalian systems alone. Collectively, our findings provide a rationale for co-targeting MEK and GPCR-Gαq-PLCβ-PKC signaling as a strategy to overcome acquired resistance and extend the durability of MAPK-directed therapies in patients with MAPK-driven cancers.

## Materials and methods

### Worm strains and culture

*C.elegans* were maintained according to the standard methods (Brenner 1974). All worms were cultivated at 20 °C on nematode growth medium (NGM) plates seeded with the *Escherichia coli* OP50 unless stated otherwise. Adult hermaphrodite worms were used in the live-cell imaging experiments. The wild-type strain was Bristol N2. Some strains were provided by the Caenorhabditis Genetics Center (CGC), which is funded by the NIH Office of Research Infrastructure Programs (P40 OD010440). Table S1 summarizes the strains used in this study.

### Genetic screens

We used forward genetic screens to isolate*lin-45(V627E)* suppressors. The*lin-45 (V627E)*mutant animals (P0) were synchronized at the late L4 larval stage, collected in 4 mL M9 buffer, and incubated with 50 mM ethyl methanesulfonate (EMS) for 4 hours at room temperature with constant rotation. Animals were then washed with M9 for three times and cultured under the standard conditions. After 20 hours, adult animals were bleached. Eggs (F1) were distributed and raised on ∼800 9-cm NGM plates, each containing 50 to 100 eggs. Adult F2 animals on each plate were screened. Mutant animals with normal vulva were individually cultured, and their progenies were further observed. We identified mutations using whole-genome sequencing and mapped *egl-30(R25C)*on chromosome I. We confirmed gene cloning using multiple alleles.

### Brood Size Assay

Synchronized L4 worms were transferred singly to NGM plates with *E.coli* OP50 bacteria cultured with Luria-Bertani broth. Worms were repeatedly transferred to a freshly seeded NGM plate every 24 h until reproduction ceased. We counted the number of progeny at their L3/L4 stages. Our measured WT brood size is consistent with the previous results. Total brood sizes were analyzed using Kruskal-Wallis test followed by Dunn’s multiple comparisons test (GraphPad Prism 10). Data are presented as mean ± SD, and p < 0.05 was considered statistically significant. Each data point represents an individual worm, with experiments repeated independently at least three times.

### Cell culture and treatment

The human melanoma cell line A375 was a gift from Prof. Jiang Peng (Tsinghua University) and was cultured in DMEM (Gibco) containing 10 % fetal bovine serum (Gibco) and 1 % penicillin and streptomycin (Yeasen) at 37 ℃ with 5 % CO_2_. A375-TR (A375-trametinib resistant) cell lines were established by culturing parental A375 cells in increasing doses of MEKi trametinib, followed by isolation of individual clones. Resistant lines were maintained in trametinib (10 nM) > 6 months (Villanueva, et al. 2010).

Short hairpin RNA (shRNA) constructs targeting human *GNA11*, *GNAQ*and *F2RL1* were designed and generated via molecular cloning. Five target sequences were used: shGαq/11#1, 5′-CCGGAGCTCAAGCTGCTGCTGCTCGCTCGAGCGAGCAGCAGCAGCTTGAGCTTTTTTG-3′, shGαq/11#2, 5′-CCGGACCCCTGGTTCCAGAACTCCTCTCGAGAGGAGTTCTGGAACCAGGGGTTTTTTG-3′, shF2RL1#1, 5′-CCGGCGAAACCTCATCTCTACTAAACTCGAGTTTAGTAGAGATGAGGTTTCGTTTTTG-3′, shF2RL1#2, 5′-CCGGGCTCTTTGTAATGTGCTTATTCTCGAGAATAAGCACATTACAAAGAGCTTTTTG-3′, and shF2RL1#3, 5′-CCGGCCACTGTTAAGACCTCCTATTCTCGAGAATAGGAGGTCTTAACAGTGGTTTTTG-3′. A corresponding empty vector was purchased from Thermo Fisher Scientific and used as a control (Ctrl). Lentiviral particles were produced and used to infect A375 and A375-TR melanoma cells. Stable cell lines expressing shRNA were selected according to the manufacturer’s instructions using puromycin and maintained under selection for downstream experiments.

### Cell viability assays and combination index calculation

The Cell-ATP Viability Detection Kit (HY-K0302; MedChemExpress) was adopted to determine the Cell viability of each group. Cells were incubated with serial dilutions of drugs alone or two-drug combinations in complete culture medium for 72 h. At least three independent experiments were performed. The curve-fitting software, GraphPad Prism version 10 (GraphPad Software, SanDiego, CA, USA), was used to calculate half-maximal inhibitory concentration values. Combination index (CI) values were calculated using the Chou-Talalay method (Chou and Talalay 1984) implemented in the synergy Python package (Wooten and Albert 2021).

### Clonogenic assay

Cells were seeded in six-well plates at low density (100 cells/well) and cultured with drugs for 10 days, after which colonies were fixed with methanol and stained with 1× crystal violet. The quantification of the average colony area were analysed by using ImageJ.

### Mouse xenografts and in vivo drug studies

All animal experiments were approved by the Animal Care and Use Committee of Tsinghua University. Female BALB/c-nude mice (5 weeks old) were housed under specific pathogen-free conditions in a temperature-controlled environment (approximately 21 °C) with a 12 h light/12 h dark cycle and ad libitum access to food and water.

Parental A375 or trametinib-resistant A375 (A375-TR) melanoma cells (5 × 10^6^ cells) were resuspended in PBS and injected subcutaneously into the flanks of mice ( ≥ 5 mice per group). Tumor growth was monitored twice weekly using caliper measurements, and tumor volume was calculated using the modified ellipsoid formula: volume = 1/2 × (length × width^2^). When tumors reached approximately 250 mm³, mice were randomized into four treatment groups: vehicle control, trametinib alone, PLCβ/PKC pathway inhibitor alone (U73122 or sotrastaurin), or the combination of trametinib with U73122 or sotrastaurin.

Trametinib (0.2 mg/kg; #C16249920, Macklin) and sotrastaurin (80 mg/kg; #HY-10343, MCE) were formulated in vehicle (5% DMSO, 40% PEG300, 5% Tween-80, and 50% sterile saline, v/v) and administered by oral gavage at a volume of 100 μL per mouse. U73122 (50 mg/kg; #HY-13419, MCE) was ground into fine powder, suspended in sterile saline, and administered by intraperitoneal injection (100 μL per mouse). Treatments were generally performed three times per week. Based on tolerability assessments indicating body weight loss with more frequent dosing, U73122 was administered three times per week in parental A375 xenografts and twice per week in A375-TR xenografts. Control animals received vehicle or sterile saline using the corresponding administration routes and schedules. Mouse body weight and general condition were monitored throughout the treatment period.

### Western Blot Analysis

Cells were washed three times with ice-cold phosphate-buffered saline (PBS) to remove residual medium and debris, and lysed in RIPA buffer (#C0121, Beyotime) supplemented with complete protease inhibitors and PhosSTOP phosphatase inhibitors (Roche). Lysates were clarified by centrifugation at 14,000 × g for 15 min at 4 °C, and supernatants were collected and stored at −80 °C until use. Protein samples were mixed with 5 × SDS loading buffer (#20315ES20, Yeasen), boiled at 95 °C for 5 min, and resolved by SDS-PAGE using FastPAGE precast gels (#TSP024-15, TSINGKE). Proteins were transferred onto PVDF membranes (#B36011, ESAYBIO), which were blocked in 5% bovine serum albumin (BSA) and incubated with primary antibodies overnight at 4 °C.

After washing three times with TBST (TBS containing 0.1% Tween-20; #T1081, Solarbio), membranes were incubated with HRP-conjugated secondary antibodies for 1 h at room temperature. Immunoreactive bands were detected using SuperSignal West Dura chemiluminescent substrate (Thermo Fisher Scientific, #34076).

The following antibodies were used: anti-α-tubulin (DM1A; #3873T, CST), anti-Gαq (D5V1B; #14373S, CST), anti-GNA11 (ActivAb™ Anti-GNA11 polyclonal antibody; #K003471P, Solarbio), anti-PAR2 (D61D5; #6976, CST), anti-ERK1/2 (p44/42 MAPK; #4695T, CST), and anti-phospho-ERK1/2 (Thr202/Tyr204; #4370T, CST). HRP-linked anti-mouse (#7076P2, CST) and anti-rabbit (#7074P2, CST) IgG antibodies were used as secondary antibodies.

### RNA sequencing

Synchronized worms were cultured on NGM plates with the OP50 bacteria. Day 1 adult worms were harvested 24 hours after the fourth larval (L4) stage and then lysed with TRIzol reagent (Invitrogen). For cell lines, after treatment with serial dilutions of drugs in complete culture medium for 24 h, A375 or A375-TR cells were lysed with TRIzol reagent (Invitrogen). Total RNA was extracted according to the manufacturer’s protocol. SUPERase•InTM RNase Inhibitor (Invitrogen) was used in each step to prevent RNA from degradation. RNA concentration was quantified using the Qubit RNA High Sensitivity Assay Kit (Invitrogen). RNA quality was assessed with the Agilent 2100 bioanalyzer system, and samples with an RNA integrity number (RIN) above 6.0 were used to construct the library, 50 ng to 500 ng of total RNA was used for library preparation using the KAPA RNA HyperPrep Kit (KAPA Biosystems). Library samples were analyzed by Agilent 2100 bioanalyzer system and Qubit for quality control and quantification. The samples were sequenced on an Illumina HiSeq platform. A total of 150-bp paired-end reads were generated.

### Gene expression analysis

The resulting raw reads were assessed for quality, adaptor content, and duplication rates with FastQC. The raw sequencing reads were trimmed using Trim_galore (version 0.4.4) to remove the low-quality bases and adaptor sequences. Paired-end reads with at least 20 nucleotides in length were aligned to *C. elegans* reference genome (ce10) using STAR (2.5.4b) with the parameter ‘-sjdbOverhang 139’. The numbers of reads that aligned to genes were quantified by HTSeq (version 0.9.1). Only uniquely mapped reads were used to calculate the relative expression level of the gene. Reads overlapping with multiple genes or aligning to multiple regions were excluded. Gene names were annotated on the DAVID website (https://david.ncifcrf.gov/). Differentially expressed genes were identified using DESeq2 package in R programming language, and differentially expressed genes were defined with the following criteria: up-regulated genes (false discovery rate (FDR) less than 0.05, log2 fold change greater than 1, un-normalized counts from HTSeq greater than 3); downregulated genes (FDR less than 0.05, log2 fold change less than −1, and un-normalized counts from HTSeq greater than 3); and other genes representing the ones that are not differentially expressed. We used the ggplot2 package of R to plot figures.

### Molecular biology

Short hairpin RNAs (shRNAs) targeting *GNA11,GNAQ* and *F2RL1* were designed based on published sequences. Oligonucleotides encoding the hairpin sequences were synthesized, annealed, and cloned into the vector pLKO.1 plasmid downstream of the U6 promoter via standard molecular cloning techniques following standard protocols. Ligated products were transformed into *E. coli* DH5α, and positive clones were confirmed by colony PCR and Sanger sequencing. All the plasmids and primers were listed in Table S2.

### Imaging

Adult *C.elegans* hermaphrodites were anesthetized using 1 μM levamisole in M9 buffer, mounted on 3% agarose pads, maintained at room temperature and imaged immediately. A Zeiss LSM900 confocal microscope equipped with a 100×/1.46 oil objective and controlled by ZEN software (Carl Zeiss) was utilized. All images were acquired using a Zeiss Axio Observer (bright-field mode).

## Data availability

Data and materials availability: All data are available in the main text or supplementary materials. All sequencing data from this study have been submitted to the NCBI Sequence Read Archive (SRA; www.ncbi.nlm.nih.gov/sra) with BioProject accession numbers PRJNA1404971 and PRJNA1405274.

## Acknowledgements

We thank Prof. Stephan Vagner and Prof. Peng Jiang for valuable discussions and insightful comments on the manuscript. This work was supported by the National Key R&D Program of China (Grant Nos. 2022YFA1302700 and 2024YFA1307301) and the National Natural Science Foundation of China (Grant Nos. 32270773 and 32470730 to W.L.; 92254306, 32430026 and 32021002 to G.O.). Additional support was provided by the TsienTang Life Science Development Fund at Tsinghua University.

## Author contributions

G.O. and W.L. designed research; J.K. and Z.W. performed research; J.K. and Z.G. analyzed data; J.K. and W.L. wrote the paper.

