## Supplemental Figure 1, 2, 3, 4, 5, 6, 7, Supplemental Table 1, 2 for "GPCR-Gαq Signaling Drives Adaptive Resistance to MEK Inhibition in BRAF-Mutant Melanoma"

### Figure S1

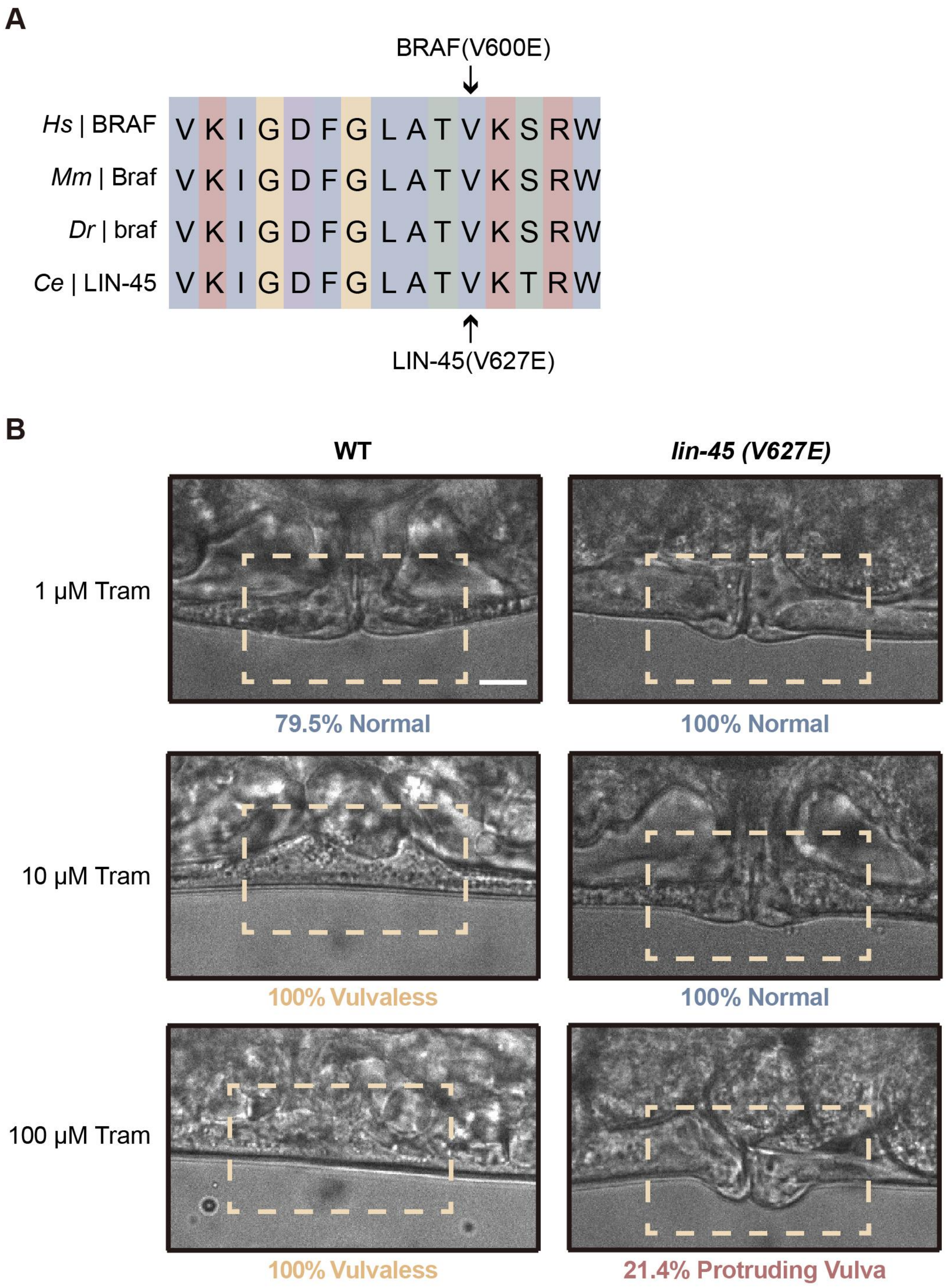
A
BRAF(V600E)
Hs | BRAF VKIGDFGLATVKSRW
Mm | Braf VKIGDFGLATVKSRW
Dr | braf VKIGDFGLATVKSRW
Ce | LIN-45 VKIGDFGLATVKTRW
LIN-45(V627E)
B
WT
lin-45 (V627E)
1 μM Tram
79.5% Normal
100% Normal
10 μM Tram
100% Vulvaless
100% Normal
100 μM Tram
100% Vulvaless
21.4% Protruding Vulva

**Figure S1. Conservation and functional characterization of the LIN-45(V627E) mutation in *C. elegans*.**

**(A)** Multiple sequence alignment of *C. elegans* LIN-45 and its homologs in human (*Homo sapiens, Hs*), mouse (*Mus musculus, Mm*), and zebrafish (*Danio rerio, Dr*). The arrow indicates the conserved valine residue in the activation loop, whose substitution by glutamate (V627E) mimics phosphorylation and confers constitutive kinase activation.

**(B)** Representative bright-field images of vulval morphology in wild-type and *lin-45(V627E)* animals treated with DMSO or trametinib (Tram) at the indicated concentrations. n > 30 animals per condition. Scale bar, 100 µm.

### Figure S2

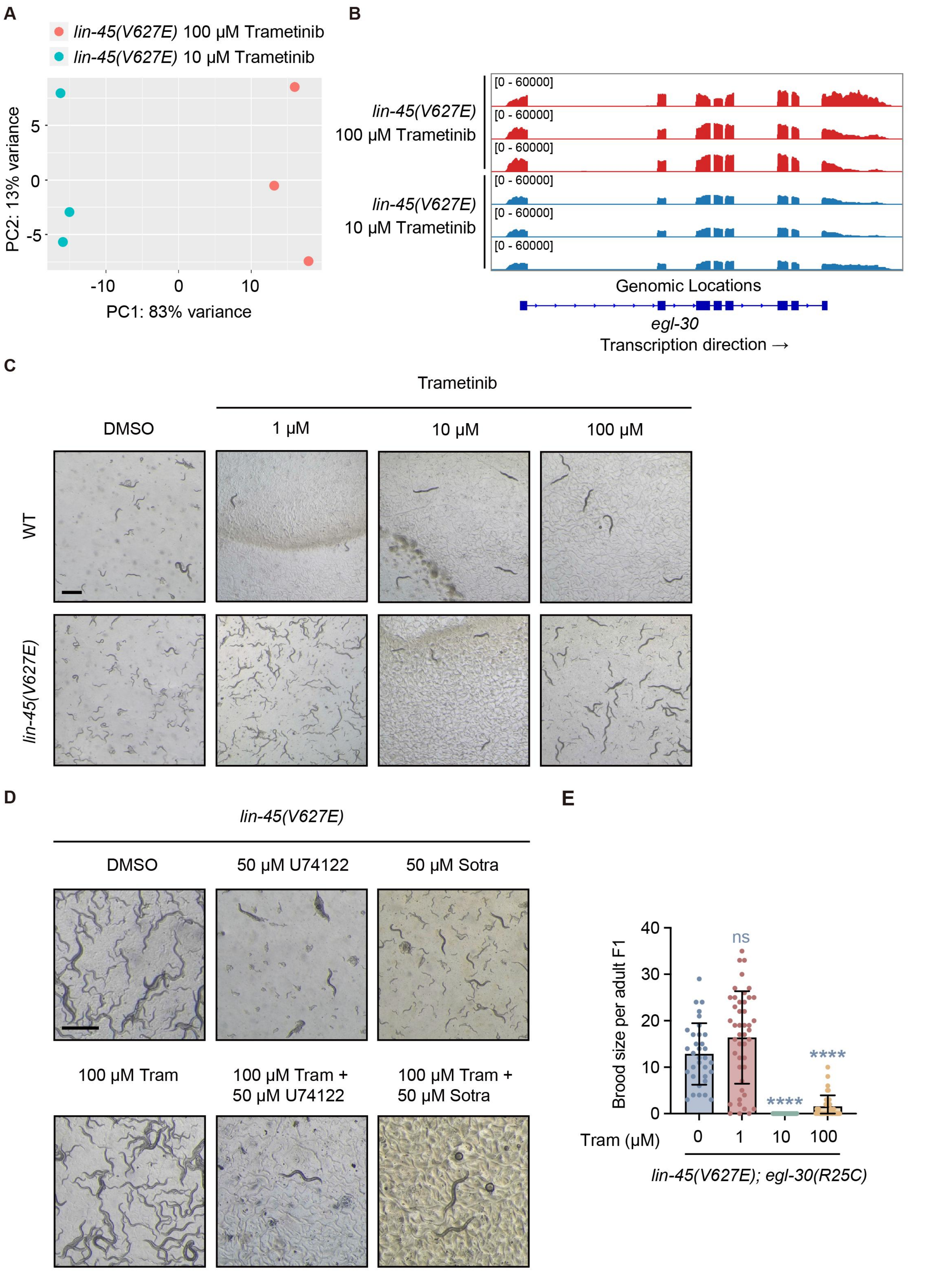
A
lin-45(V627E) 100 μM Trametinib
lin-45(V627E) 10 μM Trametinib
PC2: 13% variance
PC1: 83% variance
B
lin-45(V627E) 100 μM Trametinib
lin-45(V627E) 10 μM Trametinib
[0 - 60000]
Genomic Locations
egl-30
Transcription direction →
C
Trametinib
DMSO
1 μM
10 μM
100 μM
WT
lin-45(V627E)
D
lin-45(V627E)
DMSO
50 μM U74122
50 μM Sotra
100 μM Tram
100 μM Tram + 50 μM U74122
100 μM Tram + 50 μM Sotra
E
Brood size per adult F1
ns
****
****
Tram (μM)
0
1
10
100
lin-45(V627E); egl-30(R25C)

**Figure S2. Upregulation of *egl-30*/Gαq and requirement of Gαq-PLCβ-PKC signaling for acquired trametinib resistance in *lin-45(V627E)* mutants.**

**(A)** Principal component analysis (PCA) of RNA-seq profiles from *lin-45(V627E)* F1 animals treated with 100 μM or 10 μM trametinib (n = 3 independent biological replicates per condition).

**(B)** Integrative Genomics Viewer (IGV) tracks showing normalized RNA-seq read coverage at the *egl-30* locus in *lin-45(V627E)* F1 animals treated with 100 μM or 10 μM trametinib.

**(C-D)** Representative bright-field images of wild-type (WT) and *lin-45(V627E)* animals, along with their progeny, following treatment with DMSO, trametinib (Tram) at the indicated concentrations, U73122 (50 μM), Sotrastaurin (50 μM), or the indicated drug combinations for 10 days. Five adult animals were transferred per plate. Scale bar, 1 mm.

**(E)** Dose-dependent effects of trametinib (Tram) on brood size in *lin-45(V627E); egl-30(R25C)* F1 animals. Statistical analysis was performed using Kruskal-Wallis test followed by Dunn's multiple comparisons test. n > 30 animals per condition. ****P < 0.0001; ns, not significant.

### Figure S3

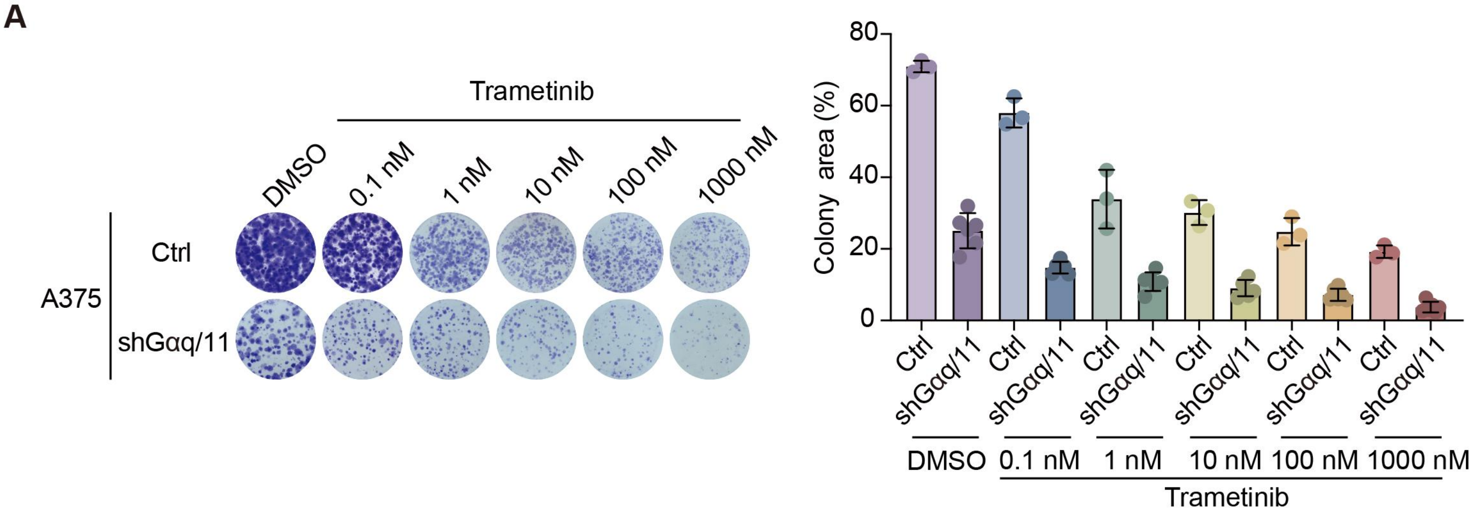

A
Trametinib
DMSO
0.1 nM
1 nM
10 nM
100 nM
1000 nM
A375
Ctrl
shGαq/11
Colony area (%)
80
60
40
20
0
Ctrl
shGαq/11
DMSO
0.1 nM
1 nM
10 nM
100 nM
1000 nM
Trametinib


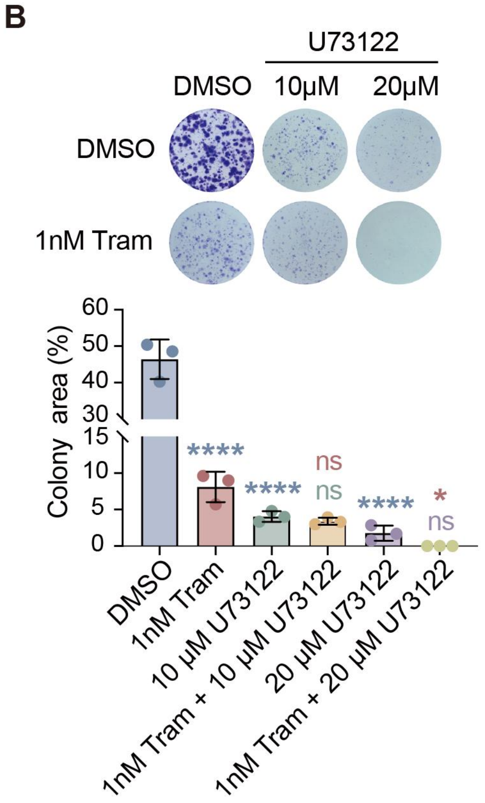

B
U73122
DMSO
10μM
20μM
DMSO
1nM Tram
Colony area (%)
60
50
40
30
15
10
5
0
****
****
ns
ns
****
*
ns
DMSO
1nM Tram
10 μM U73122
1nM Tram + 10 μM U73122
20 μM U73122
1nM Tram + 20 μM U73122


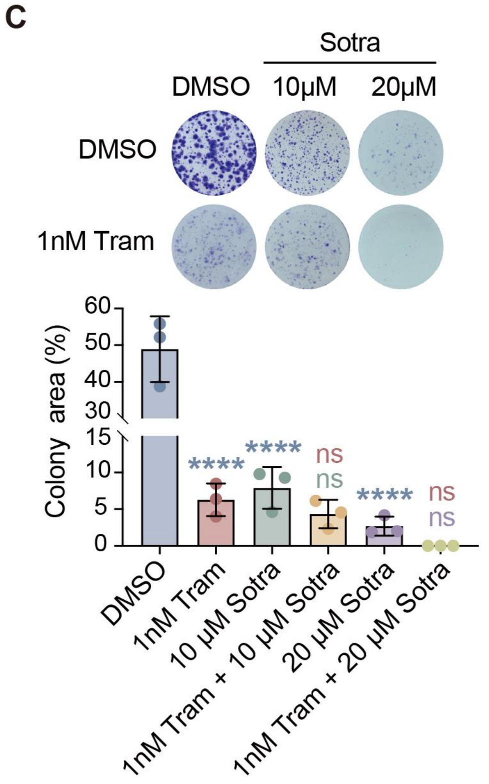

C
Sotra
DMSO
10μM
20μM
DMSO
1nM Tram
Colony area (%)
60
50
40
30
15
10
5
0
****
****
ns
ns
****
ns
ns
DMSO
1nM Tram
10 μM Sotra
1nM Tram + 10 μM Sotra
20 μM Sotra
1nM Tram + 20 μM Sotra

**Figure S3. Inhibition of Gαq-PLCβ-PKC signaling reduces clonogenic growth of parental A375 cells.**

**(A)** Colony formation assay of A375 cells transduced with control vector (Ctrl) or shGαq/11 and treated with vehicle or trametinib at the indicated concentrations. Representative images of crystal violet-stained colonies (left) and quantification of colony area (right) are shown. n = 3 independent biological replicates.

**(B)** Colony formation assay of A375 cells treated with vehicle, trametinib (1 nM), U73122 (10 μM or 20 μM), or the indicated combinations. Representative images (top) and quantification of colony area (bottom) are shown. *P < 0.1, ****P < 0.0001; ns, not significant; one-way ANOVA with Tukey's multiple comparisons test. n = 3 independent biological replicates.

**(C)** Colony formation assay of A375 cells treated with vehicle, trametinib (1 nM), sotrastaurin (10 μM or 20 μM), or the indicated combinations. Representative images (top) and quantification of colony area (bottom) are shown. ****P < 0.0001; ns, not significant; one-way ANOVA with Tukey's multiple comparisons test. n = 3 independent biological replicates.

### Figure S4

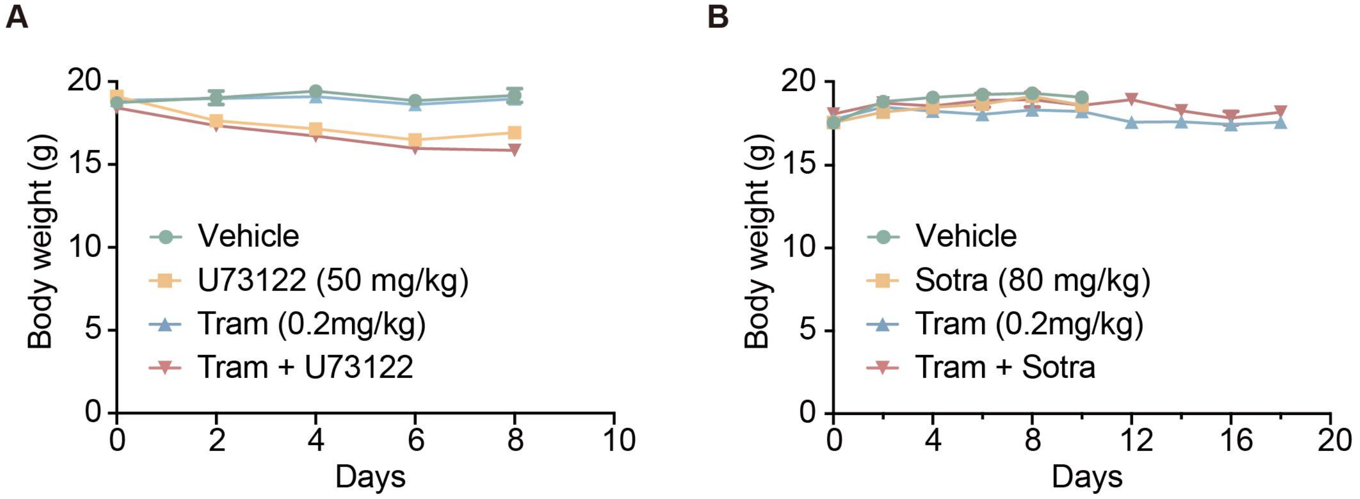
A
B
Body weight (g)
Days
Vehicle
U73122 (50 mg/kg)
Tram (0.2mg/kg)
Tram + U73122
Sotra (80 mg/kg)
Tram + Sotra

**Figure S4. Body weight monitoring of A375 xenograft-bearing mice during treatment.**

**(A)** Body weights of parental A375 xenograft-bearing mice treated with vehicle, U73122 (50 mg/kg), trametinib (0.2 mg/kg), or their combination (n = 17 mice per group).

**(B)** Body weights of parental A375 xenograft-bearing mice treated with vehicle, sotrastaurin (Sotra; 80 mg/kg), trametinib (0.2 mg/kg), or their combination (n = 12 mice per group).

Data represent mean ± SEM. Treatments were initiated when mean tumor volume reached ~250 $mm^3$.

### Figure S5

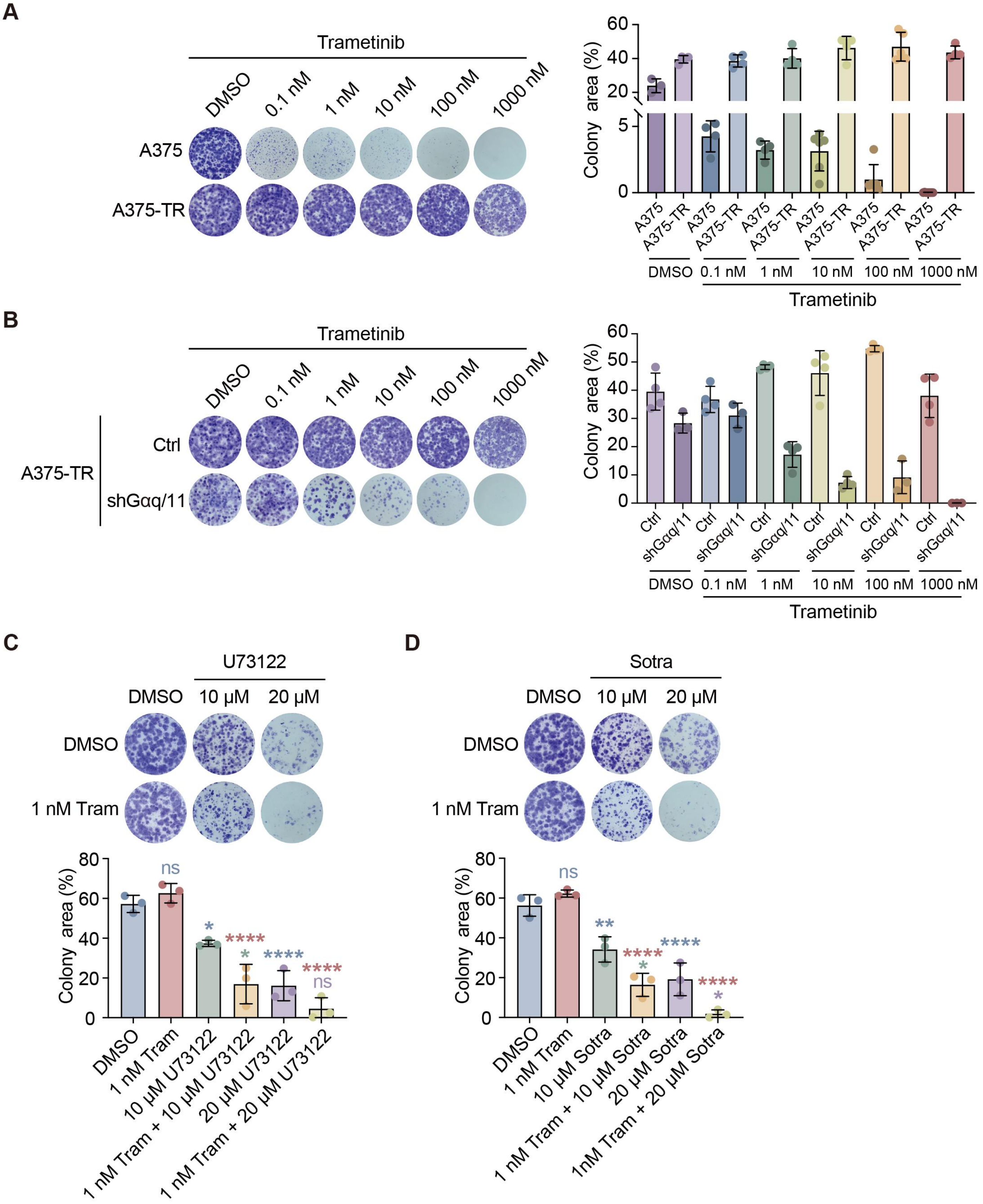
A
Trametinib
DMSO
0.1 nM
1 nM
10 nM
100 nM
1000 nM
A375
A375-TR
Colony area (%)
A375
A375-TR
DMSO
0.1 nM
1 nM
10 nM
100 nM
1000 nM
Trametinib
B
Trametinib
A375-TR
Ctrl
shGαq/11
Colony area (%)
DMSO
0.1 nM
1 nM
10 nM
100 nM
1000 nM
Trametinib
C
U73122
DMSO
10 μM
20 μM
DMSO
1 nM Tram
Colony area (%)
ns
*
****
*
****
****
ns
DMSO
1 nM Tram
10 μM U73122
1 nM Tram + 10 μM U73122
20 μM U73122
1 nM Tram + 20 μM U73122
D
Sotra
DMSO
10 μM
20 μM
DMSO
1 nM Tram
Colony area (%)
ns
**
****
*
****
****
*
DMSO
1 nM Tram
10 μM Sotra
1 nM Tram + 10 μM Sotra
20 μM Sotra
1nM Tram + 20 μM Sotra

**Figure S5. Inhibition of Gαq-PLCβ-PKC signaling reduces clonogenic growth of trametinib-resistant A375 cells.**

**(A)** Colony formation assay of parental A375 and trametinib-resistant A375 (A375-TR) cells treated with vehicle or trametinib at the indicated concentrations. Representative images of crystal violet-stained colonies (left) and quantification of colony area (right) are shown. n = 3 independent biological replicates.

**(B)** Colony formation assay of A375-TR cells transduced with control vector (Ctrl) or shGαq/11 and treated with vehicle or trametinib at the indicated concentrations. Representative images (left) and quantification of colony area (right) are shown. n = 3 independent biological replicates.

**(C)** Colony formation assay of A375-TR cells treated with vehicle, trametinib (1 nM), U73122 (10 μM or 20 μM), or the indicated combinations. Representative images (top) and quantification of colony area (bottom) are shown. *P < 0.1, ****P < 0.0001; ns, not significant; one-way ANOVA with Tukey's multiple comparisons test. n = 3 independent biological replicates.

**(D)** Colony formation assay of A375-TR cells treated with vehicle, trametinib (1 nM), sotrastaurin (10 μM or 20 μM), or the indicated combinations. Representative images (top) and quantification of colony area (bottom) are shown. *P < 0.1, **P < 0.01, ****P < 0.0001; ns, not significant; one-way ANOVA with Tukey's multiple comparisons test. n = 3 independent biological replicates.

### Figure S6

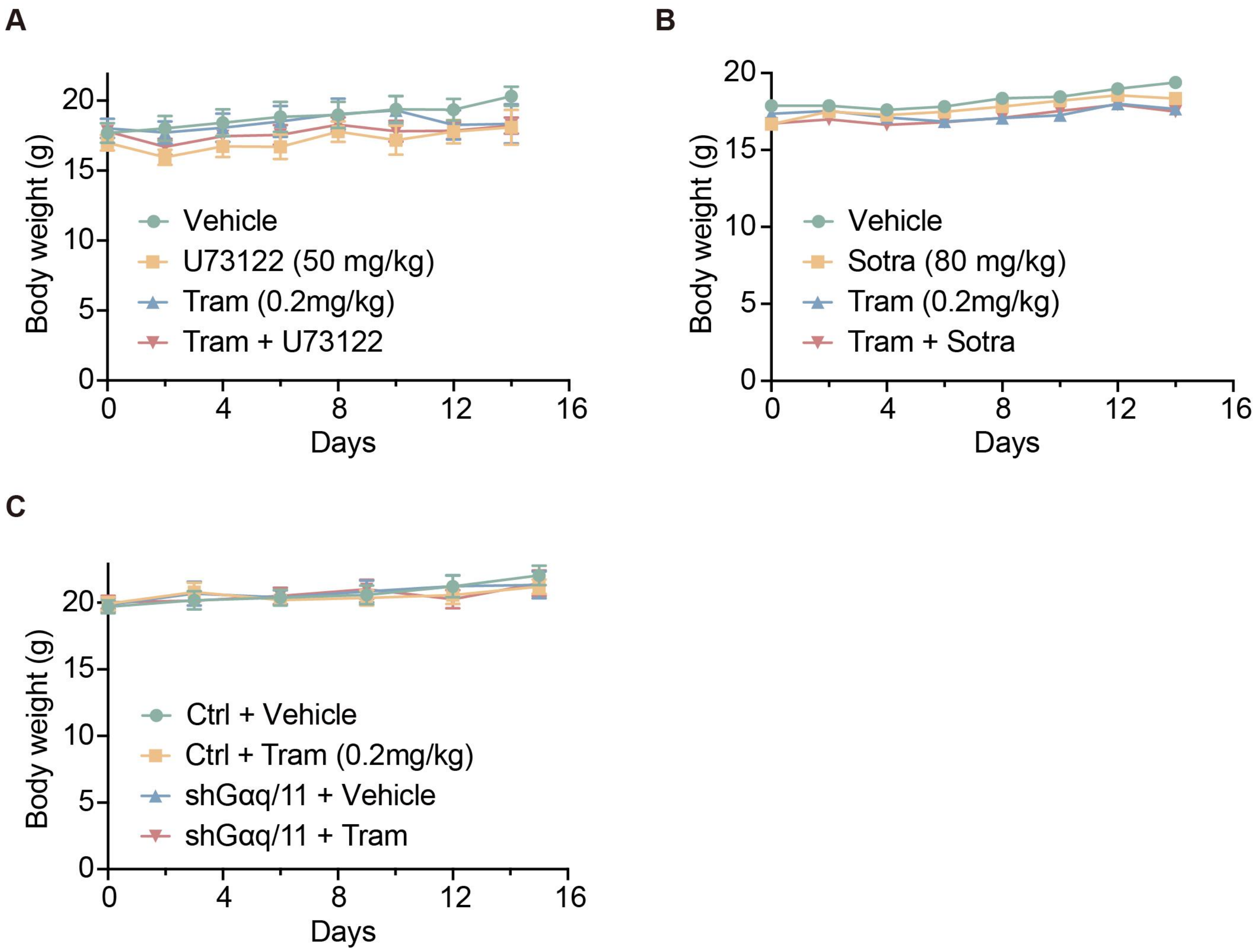
A
Body weight (g)
Days
Vehicle
U73122 (50 mg/kg)
Tram (0.2mg/kg)
Tram + U73122
B
Body weight (g)
Days
Vehicle
Sotra (80 mg/kg)
Tram (0.2mg/kg)
Tram + Sotra
C
Body weight (g)
Days
Ctrl + Vehicle
Ctrl + Tram (0.2mg/kg)
shGαq/11 + Vehicle
shGαq/11 + Tram

**Figure S6. Body weight monitoring of A375-TR xenograft-bearing mice during treatment.**

**(A)** Body weights of trametinib-resistant A375-TR xenograft-bearing mice treated with vehicle, U73122 (50 mg/kg), trametinib (0.2 mg/kg), or their combination (n = 10 mice per group).

**(B)** Body weights of A375-TR xenograft-bearing mice treated with vehicle, sotrastaurin (80 mg/kg), trametinib (0.2 mg/kg), or their combination (n = 10 mice per group).

**(C)** Body weights of mice bearing A375-TR xenografts transduced with control vector (Ctrl) or shGαq/11 and treated with vehicle or trametinib (0.2 mg/kg) (n = 18 mice per group).

Data represent mean ± SEM. Treatments were initiated when mean tumor volume reached ~250 $mm^3$.

#### Figure S7

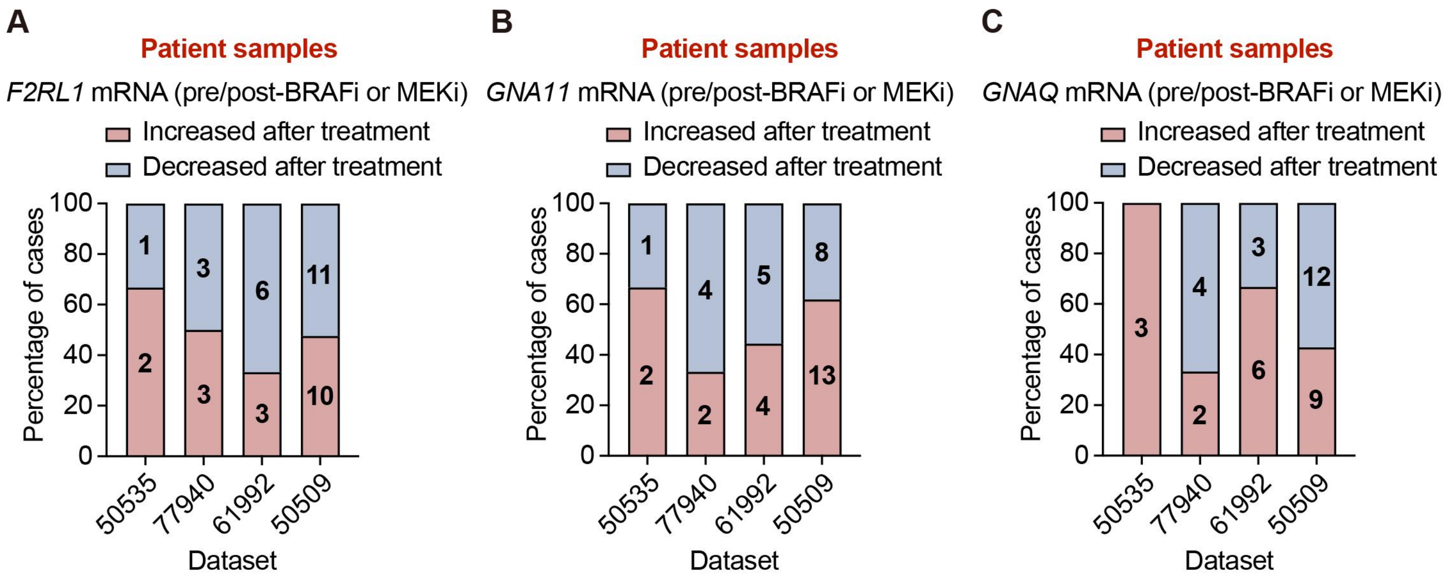

A
Patient samples
F2RL1 mRNA (pre/post-BRAFi or MEKi)
Increased after treatment
Decreased after treatment
Percentage of cases
0
20
40
60
80
100
1
2
3
3
6
3
11
10
50535
77940
61992
50509
Dataset
B
Patient samples
GNA11 mRNA (pre/post-BRAFi or MEKi)
Increased after treatment
Decreased after treatment
Percentage of cases
1
2
4
2
5
4
8
13
50535
77940
61992
50509
Dataset
C
Patient samples
GNAQ mRNA (pre/post-BRAFi or MEKi)
Increased after treatment
Decreased after treatment
Percentage of cases
3
4
2
3
6
12
9
50535
77940
61992
50509
Dataset

**Figure S7. Dataset-specific expression changes of *F2RL1* (PAR2), *GNAQ*, and *GNA11* in melanoma samples before and after MAPK pathway inhibitor treatment.**

**(A-C)** Paired human melanoma samples from individual datasets (RNA-seq: 50535, 77940, 61992; microarray: 50509) were analyzed for *F2RL1* (PAR2), *GNAQ*, and *GNA11* mRNA levels pre-/post-treatment with BRAFi, MEKi, or combined BRAFi/MEKi. Numbers in each column indicate the number of cases.

**Table S1.** ***Caenorhabditis elegans*** **strains used in this study.**

| Strain Name | Genotype | Source / Generation |
|---|---|---|
| N2 | Wild type | Bristol N2 reference strain. |
| PHX4962 | *lin-45 (syb4962) IV.* | Purchased from SunyBiotech Co., Ltd. Strain generated by CRISPR-Cas9-mediated genome editing. |
| MT1434 | *egl-30 (n686) I.* | *Caenorhabditis* Genetics Center (CGC). |
| DA1084 | *egl-30 (ad806) I.* | |
| GOU4414 | *lin-45 (syb4962) IV; egl-30 (cas30009) I.* | Generated by EMS mutagenesis in this study. |
| GOU4415 | *lin-45 (syb4962) IV; egl-30 (n686) I.* | Generated by genetic crossing in this study. |
| GOU4416 | *lin-45 (syb4962) IV; egl-30 (ad806) I.* | |

Table S2. Plasmids and primers used in this study.

| Plasmid | Forward Primer | Reverse Primer | Notes |
|---|---|---|---|
| pLKO.1-puro Control Vector (SHC001) | - | - | Lentiviral shRNA backbone control vector. Obtained from the Tsinghua University shRNA Library; original construct sourced from Sigma-Aldrich (MISSION® pLKO.1-puro SHC001). |
| pLKO.1-shGαq/11#1 | GCTCGCTCGAGCGAGCAGCAGCAGCTTGAGCTTTTTTGATTCTCGACCTCGA | GCTCGCTCGAGCGAGCAGCAGCAGCTTGAGCTCCGGTGTTTCGTCCTTTCCA | shRNA targeting *GNAQ/GNA11* cloned into pLKO.1-puro backbone by PCR amplification from SHC001 vector. |
| pLKO.1-shGαq/11#2 | CTATTCTCGAGAATAGGAGGTCTTAACAGTGGTTTTTGATTCTCGACCTCGA | CTATTCTCGAGAATAGGAGGTCTTAACAGTGGCCGGTGTTTCGTCCTTTCCA | shRNA targeting *GNAQ/GNA11* cloned into pLKO.1-puro backbone by PCR amplification from SHC001 vector. |
| pLKO.1-shF2RL1#1 | CTAAACTCGAGTTTAGTAGAGATGAGGTTTCGTTTTTGATTCTCGACCTCGA | CTAAACTCGAGTTTAGTAGAGATGAGGTTTCGCCGGTGTTTCGTCCTTTCCA | shRNA targeting *F2RL1* cloned into pLKO.1-puro backbone by PCR amplification from SHC001 vector. |
| pLKO.1-shF2RL1#2 | TTATTCTCGAGAATAAGCACATTACAAAGAGCTTTTTGATTCTCGACCTCGA | TTATTCTCGAGAATAAGCACATTACAAAGAGCCCGGTGTTTCGTCCTTTCCA | shRNA targeting *F2RL1* cloned into pLKO.1-puro backbone by PCR amplification from SHC001 vector. |
| pLKO.1-shF2RL1#3 | CTATTCTCGAGAATAGGAGGTCTTAACAGTGGTTTTTGATTCTCGACCTCGA | CTATTCTCGAGAATAGGAGGTCTTAACAGTGGCCGGTGTTTCGTCCTTTCCA | shRNA targeting *F2RL1* cloned into pLKO.1-puro backbone by PCR amplification from SHC001 vector. |
